# Salience network atrophy links neuron type-specific pathobiology to loss of empathy in frontotemporal dementia

**DOI:** 10.1101/691212

**Authors:** Lorenzo Pasquini, Alissa L. Nana, Gianina Toller, Jesse Brown, Jersey Deng, Adam Staffaroni, Eun-Joo Kim, Ji-Hye L. Hwang, Libo Li, Youngsoon Park, Stephanie E. Gaus, Isabel Allen, Virginia E. Sturm, Salvatore Spina, Lea T. Grinberg, Katherine P. Rankin, Joel Kramer, Howard H. Rosen, Bruce L. Miller, William W. Seeley

**Author notes:** Corresponding author: William W. Seeley, MD. 675 Nelson Rising Lane 94158, San Francisco, California USA.

## Abstract

Each neurodegenerative syndrome reflects a stereotyped pattern of cellular, regional, and large-scale brain network degeneration. In behavioral variant frontotemporal dementia (bvFTD), a disorder of social-emotional function, von Economo neurons (VENs) and fork cells are among the initial neuronal targets. These large layer 5 projection neurons are concentrated in the anterior cingulate and frontoinsular (FI) cortices, regions that anchor the salience network, a large-scale system linked to social-emotional function. Here, we studied patients with bvFTD, amyotrophic lateral sclerosis (ALS), or both, given that these syndromes share common pathobiological and genetic factors. Our goal was to determine how neuron type-specific TAR DNA-binding protein of 43 kDa (TDP-43) pathobiology relates to atrophy in specific brain structures and to loss of emotional empathy, a cardinal feature of bvFTD. We combined questionnaire-based empathy assessments, *in vivo* structural MR imaging, and quantitative histopathological data from 16 patients across the bvFTD/ALS spectrum. We show that TDP-43 pathobiology within right FI VENs and fork cells is associated with salience network atrophy spanning insular, medial frontal, and thalamic regions. Gray matter degeneration within these structures mediated loss of emotional empathy, suggesting a chain of influence linking the cellular, regional/network, and behavioral levels in producing signature bvFTD clinical features.

## Introduction

Growing evidence suggests that misfolded neurodegenerative disease proteins accumulate within vulnerable onset regions, presumably within the most vulnerable neurons therein, before spreading to other brain areas along large-scale brain network connections (Brettschneider et al. 2015; Seeley 2017). Studies in behavioral variant frontotemporal dementia (bvFTD) have revealed that von Economo neurons (VENs) and fork cells are the likely initial targets (Kim et al. 2012; Seeley et al. 2012; Santillo et al. 2013; Nana et al. 2018). These neuronal morphotypes make up a unique class of large, bipolar layer 5 projection neurons, which, in large-brained highly social mammals, are concentrated in the anterior cingulate and anterior agranular insular (i.e. frontoinsular [FI]) cortices (Nimchinsky et al. 1995; Allman et al. 2011; Stimpson et al. 2011). The anterior cingulate cortex and FI are known for their functional coactivation as part of the salience network, a large-scale brain system associated with homeostatic behavioral guidance (Seeley et al. 2007, 2012; Pasquini et al. 2019). Among all salience network nodes, the right FI has been proposed to play a central role as a major cortical hub for representing interoceptive sensory information that is critical for social-emotional processes, including emotional empathy (Craig 2009; Critchley and Harrison 2013; Leigh et al. 2013; Sturm et al. 2018).

Converging evidence suggests that disease proteins associated with bvFTD progress anatomically from each patient’s putative onset regions, which are most often in the anterior cingulate cortex or FI (Raj et al. 2012; Zhou et al. 2012; Brown et al. 2019). Accordingly, patients with bvFTD show early and progressive salience network dysfunction and degeneration that correlates with the characteristic social-emotional deficits of the syndrome (Seeley et al. 2012; Hughes et al. 2018). BvFTD is the most common syndrome within the FTD clinical spectrum and is closely linked to amyotrophic lateral sclerosis (ALS) (DeJesus-Hernandez et al. 2011; Renton et al. 2011). Individual patients may present with bvFTD, ALS, or as a blended syndrome with elements of both (FTD with motor neuron disease, bvFTD-MND). The two disorders are also united by a shared underlying biology. Pathologically, transactive response DNA binding protein of 43 kDa (TDP-43) represents the underlying disease protein in ∼50% of patients with bvFTD and nearly all with sporadic ALS (MacKenzie et al. 2010). TDP-43 is a DNA/RNA binding protein normally expressed in healthy neurons, which becomes mislocalized from the nucleus to the cytoplasm, where it aggregates into neuronal cytoplasmic inclusions (Neumann et al. 2006). FTD and ALS are usually sporadic, but they share several genetic mutations in common, the most prevalent being a hexanucleotide repeat expansion in a noncoding region of chromosome 9 open reading frame 72 (*C9orf72*) (Renton et al. 2011). Recently, we completed a quantitative neuropathological study of patients representing all points along the ALS/bvFTD continuum linked to TDP-43 proteinopathy (Nana et al. 2018). We found that early bvFTD was accompanied by disproportionate TDP-43 aggregation within VENs and fork cells. At the level of individual neurons, TDP-43 aggregation (and accompanying loss of nuclear TDP-43) was associated with striking nuclear and somatodendritic atrophy. The proportion of VENs and fork cells with TDP-43 inclusions correlated with loss of emotional empathy, as assessed by patient caregivers during life. Importantly, a minority of VENs and fork cells lacked detectable nuclear TDP-43 despite the apparent absence of a cytoplasmic inclusion; this phenomenon was most striking among *C9orf72* expansion carriers. These nuclear TDP-43 depleted cells developed morphological changes comparable to TDP-43 inclusion-bearing cells (Nana et al. 2018).

In this work, we aimed to extend previous studies by assessing the link between the cellular, large-scale network, and behavioral levels through a unique dataset that combined quantitative histopathological data, *in vivo* structural MRI, and ante-mortem questionnaire-based assessments of emotional empathy from 16 patients across the ALS/bvFTD continuum. We hypothesized that TDP-43 pathobiological changes within VENs and fork cells would predict atrophy of salience network regions, which, in turn, would relate to loss of empathic concern, the ability to prosocially respond to others’ emotions.

## Materials and Methods

### Subjects

Patients were selected from a previous histopathological study (Nana et al. 2018) based on availability of a complete neuroimaging and histopathological dataset **(Table 1**). Patients were required to have (*i*) a clinical diagnosis of bvFTD, bvFTD-MND, or ALS, *(ii*) a neuropathological diagnosis of (1) frontotemporal lobar degeneration with TDP-43 pathology (FTLD-TDP), Types B or unclassifiable, (with or without MND) or (2) ALS-TDP. We focused on FTLD-TDP Types B and unclassifiable since these are the most common forms seen in patients along the ALS/bvFTD continuum (DeJesus-Hernandez et al. 2011; Renton et al. 2011). Focusing on a single clinical and neuropathological continuum allowed us to access variation in empathy deficits, brain network atrophy patterns, and neuron type-specific pathobiology among patients. Seven patients were *C9orf72* mutation positive. Patients with disease-causing mutations other than *C9orf72* repeat expansions were not included to reduce pathological and genetic heterogeneity within the sample. Based on these criteria, we included 16 patients in total (**Table 1**): five patients with bvFTD, nine with bvFTD-MND, and two with ALS. In accordance with the declaration of Helsinki, patients or their surrogates provided written informed consent prior to participation, including consent for brain donation. The University of California San Francisco Committee on Human Research approved the study.

**Table 1.** Demographics and sample characteristics

|  |  |
| --- | --- |
| N | 16 |
| Clinical diagnoses | 2 ALS; 9 bvFTD-MND; 5 bvFTD |
| Pathological diagnoses | 4 FTLD-TDP-B; 1 FTLD-TDP-U; 6 FTLD-TDP-B, MND; 2 FTLD-TDP-U, MND; 3 ALS-TDP |
| *C9orf72* expansion carriers (-: +) | 9:7 |
| Gender (Female:Male) | 10:6 |
| Age at MRI scan in years | 60.1 (6.3) |
| Scan-death interval in years | 2.4 (1.9) |
| Handedness (Left:Ambidextrous:Right) | 0:0:16 |
| Education in years | 15.9 (2.8) |
| Total intracranial volume in liters | 1.5 (0.2) |
| Broe stage, median (range) | 1 (0-2) |
| % of TDP-43 inclusion-bearing VENs and fork cells | 16.3 (17.7) |
| % of nuclear TDP-43 depleted VENs and fork cells | 7.9 (3.0) |
| % of TDP-43 inclusion-bearing NNs | 5.3 (6.2) |
| CDR Total | 1.6 (1.3) |
| CDR Sum Of Boxes | 7.1 (5.0) |
| MMSE | 23.4 (7.8) |
| IRI, empathic concern subscale (N = 13) | 18.9 (6.7) |
| **Boston Naming Test (N = 13)** | **8.6 (8.8)** |

Unless otherwise noted, numbers in parentheses are SD of preceding mean. CDR = Clinical
Dementia Rating. IRI = Interpersonal Reactivity Index, the empathic concern subscale can reach a maximal attainable score of 28. MMSE = Mini Mental State Examination.

### Neuropsychological and emotional empathy assessment

A multidisciplinary team determined patients’ clinical diagnoses following thorough neurological, neuroimaging and neuropsychological assessments. Clinical severity was assessed using the Mini Mental State Examination (MMSE) and the Clinical Dementia Rating (CDR) scale total and sum of boxes scores, using a version of the CDR adapted for FTD (Knopman et al. 2008). In 13 out of 16 patients, the empathic concern subscale of the Interpersonal Reactivity Index (Davis 2007) was completed by the patient’s informant. The empathic concern subscale measures emotional aspects of empathy, in particular the other-centered emotional response resulting from the perception of another’s emotional state. It consist of 7 questions that can comprehensively reach a maximum score of 35. Our patient sample was screened for the presence of other neuropsychological tests related or unrelated to social-emotional function. The most broadly available test was the short form of the Boston Naming Test (Kaplan et al. 1983), available in 13 of 16 patients. In this test, the examiner presents the patient with 15 drawings of objects with decreasing word frequency, and the patient is given 20 seconds to name the depicted object. The maximal attainable score is 15. All neuropsychological assessments were obtained within 90 days of structural neuroimaging.

### Structural Neuroimaging

For nine patients, MRI scans were obtained on a Siemens Trio 3T scanner at the UCSF Neuroscience Imaging Center. A T1-weighted MP-RAGE structural scan was acquired with an acquisition time = 8 min 53 sec, sagittal orientation, a field of view of 160 × 240 × 256 mm with an isotropic voxel resolution of 1 mm3, TR = 2300 ms, TE = 2.98 ms, TI = 900 ms, flip angle = 9°. For the remaining seven patients, MRI scans were obtained on a Siemens 1.5 Tesla Magnetom scanner at the San Francisco Veterans Affairs Medical Center (SFVAMC) using a MP-RAGE sequence with a voxel resolution of 1.0 × 1.0 × 1.5 mm, TR = 10ms, TE = 4ms, TI = 300ms, flip angle = 15. Structural MRI images underwent a voxel-based morphometry analysis (Ashburner and Friston 2000) after being visually inspected for motion and scanning artifacts. Structural images were segmented in gray matter, white matter, and cerebrospinal fluid and normalized to MNI space using SPM12 (http://www.fil.ion.ucl.ac.uk/spm/software/spm12/). Gray matter images were modulated by dividing the tissue probability values by the Jacobian of the warp field, and smoothed with an isotropic Gaussian kernel with a full width at half maximum of 8 mm. The smoothed images were finally transformed to w-score maps (La Joie et al. 2012; Chételat et al. 2017). W-scores are analogous to z-scores adjusted for specific covariates (Jack et al. 1997; Boccardi et al. 2003) and reflect levels of atrophy for the specific subject after adjustment. W-score maps were derived using structural MR images from 288 cognitively normal older adults assessed at the UCSF Memory and Aging Center (**Supplementary Table S1**). These data were used to derive voxel-wise gray matter tissue intensity maps. Using this sample, multiple regression analyses were performed at the voxel-wise level to estimate gray matter intensity as adjusted for age, sex, years of education, handedness, MRI scanner, and total intracranial volume:

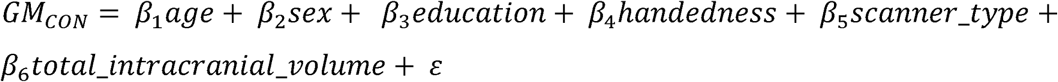

Where *GM*_*CON*_ is the voxel-specific value of a segmented gray matter tissue density map in the control sample.

Subsequently, w-score maps were computed for the patients as follows:

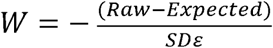

Where *Raw* is the raw value of a voxel from the smoothed image of a patient; *Expected* is the expected value for the voxel of a specific patient adjusted for covariates using the healthy control model; and *SD*ε is the standard deviation of the residuals from the healthy control model. The sign of w-score values was inverted, such that higher w-score values reflect greater atrophy. An identical approach was applied on segmented white matter tissue intensity maps to derive w-score maps of white matter atrophy. To explore the extent of gray matter atrophy in our sample, we derived a voxel-wise frequency map of gray matter atrophy using the w-score maps of all patients. Individual w-score gray matter maps were binarized at a w-score threshold of 2 and summed, such that a higher score at each voxel indicates a larger proportion of subjects with suprathreshold atrophy. Since w-score values are analogous to z-score values (La Joie et al. 2012), a w-score of 2 reflects gray matter that is over 1.96 standard deviations from the mean gray matter of healthy older adults, and as such beyond the 95% confidence interval of a normal distribution.

### Histopathology

Patients died 2.4 +/- 1.9 years after neuroimaging acquisition. Details on specimen and tissue processing can be found in previous work (Nana et al. 2018) and in **Supplementary Experimental Procedures**. The present neuropathological data represent a subset of the data reported previously (Nana et al. 2018) and no new neuropathological data were collected for the present study. Briefly, postmortem research-oriented autopsies were performed at the UCSF Neurodegenerative Disease Brain Bank. Neuropathological diagnoses were made following consensus diagnostic criteria (McKeith et al. 2005, 2017; MacKenzie et al. 2010; Montine et al. 2012; Mackenzie and Neumann 2016) based on histological and immunohistochemical methods (Tartaglia et al. 2010; Kim et al. 2012). Anatomical disease stage was assessed by a single blinded investigator (W.W.S) using an FTD rating scale (Broe et al. 2003). Blocks of the right FI were dissected, sectioned at 50 μm, and Nissl-stained to determine layer 5 within the anatomical boundaries of the right FI. Sections were stained for TDP-43 using an antibody that recognizes both full-length normal and pathological forms of the protein and counterstained with hematoxylin to allow visualization of neuron type and morphology. Finally, the numbers of VENs, fork cells, and neighboring neurons in Layer 5 with (normal) nuclear TDP-43, TDP-43 depletion, and TDP-43 inclusions were counted using the optical fractionator probe (StereoInvestigator software) to obtain the proportion of inclusion-bearing and depleted neurons of each type (Nana et al. 2018).

### Experimental Design and Statistical Analyses

In SPM12, voxel-wise regression analyses were performed on gray w-score maps using the rate of TDP-43 inclusion-bearing VENs and fork cells of the right FI as the independent variable. Models were corrected for *C9orf72* mutation status and time interval between scanning and death. A height threshold of p < 0.001 and a cluster-extend threshold of p < 0.05 FWE corrected was used in all voxel-wise analyses if not reported otherwise. This height threshold was used because a more conservative height threshold of p < 0.05 FWE corrected for multiple comparisons did not yield significant findings. The height threshold used in our study, however, is comparable to those of previous correlational neuroimaging studies on neurodegenerative disease samples of this size (Ghosh et al. 2012; Pasquini et al. 2015; Cope et al. 2020) and remains more stringent than the problematic thresholds of p < 0.01 and p < 0.05 shown to inflate false positive findings in correlative neuroimaging studies relating for example gray matter with genetic variation (Silver et al. 2011). An analogous voxel-wise model was used to assess the association between inclusion-bearing VENs and fork cells and white matter atrophy. To assess the specificity of the findings for VENs and fork cells, similar voxel-wise regression analyses were performed using the inclusion formation rate among neighboring (i.e. non-VEN, non-fork cell) layer 5 neurons in right FI as the independent variable. To explore whether the association between neuron-type specific pathobiology and structural atrophy differed between patients with and without the *C9orf72* expansion, partial residuals for average levels of atrophy were derived by regressing out the influence of time interval between scanning and death, but not *C9orf72* status, and plotted against the rate of TDP-43 inclusion-bearing VENs and fork cells separately for *C9orf72* expansion carriers and non-carriers using Matlab (https://www.mathworks.com/products/matlab.html). Secondary voxel-wise regression analyses were carried out separately on *C9orf72* mutation non-carriers. Finally, to explore the potential relevance of nuclear TDP-43 depletion (without inclusion formation) to atrophy of salience network regions, we repeated the voxel-wise gray matter regression analyses using a composite measure of TDP-43 pathobiology that summed the rate of TDP-43 inclusion-bearing and depleted VENs and fork cells of the right FI (henceforth, TDP-43 composite score). As for the inclusion-only models, partial residuals for average levels of atrophy were derived by regressing out the influence of time interval between scanning and death and plotted against the composite TDP-43 pathobiology score of VENs and fork cells. Average levels of regional gray matter atrophy associated with TDP-43 inclusion-bearing VENs and fork cells identified in the voxel-wise analyses were associated with the empathic concern score of the Interpersonal Reactivity Index using Spearman partial correlation corrected for *C9orf72* mutation status (p < 0.05 To assess the specificity of the observed associations, we performed two additional analyses using Spearman partial correlation corrected for *C9orf72* mutation status (p < 0.05): *(i)* average levels of regional gray matter atrophy associated with TDP-43 inclusion-bearing VENs and fork cells were associated with the Boston Naming Test (Kaplan et al. 1983); *(ii)* empathic concern scores were correlated with average gray matter atrophy in the right occipital cortex. For this control analysis, a region-of-interest in the right occipital cortex (region-of-interest 204; Brainnetome Atlas; https://atlas.brainnetome.org/)(Fan et al. 2016) was chosen because *(i)* this region is not part of the salience network; *(ii)* it is not typically associated with social-emotional functions; *(iii)* showed limited amounts of atrophy in our sample (**Figure 2A**); and *(iv)* was similar in size when compared to the regional atrophy clusters associated with TDP-43 inclusion-bearing VENs and fork cells (∼ 2000 voxels). Finally, mediation analysis (Baron and Kenny 1986) was conducted to test whether gray matter atrophy mediated the link between neuron type-specific pathobiology and empathy deficits. Average gray matter atrophy levels were derived from selected regions showing the strongest statistical associations with VEN/fork cell TDP-43 inclusion formation based on the voxel-wise regression analysis involving all patients. The mediation analysis was conducted using the statistical software R (https://www.r-project.org/), after scaling the measured values to z-scores.

### Data availability

The t-map derived from the voxel-wise regression analysis associating gray matter atrophy and right FI VEN and fork cell TDP-43 inclusion fraction in all patients is publicly available on NeuroVault (https://identifiers.org/neurovault.image:305261) (Gorgolewski et al. 2015). Further data that support the findings of this study are available on request from the corresponding author W.W.S. Raw data are not made publicly available because they contain information that could compromise patient privacy.

## Results

Over a span of ten years, 16 patients from across the ALS/bvFTD clinicopathological continuum met inclusion criteria (**Table 1**). Patients were assessed with antemortem structural MRI (**Figure 1A**), social-emotional function tests, and postmortem neuropathological evaluation, including quantitative assessment of TDP-43 pathobiology and neuronal densities in right FI (**Figure 1B-D**). Segmented gray matter tissue probability maps were transformed to w-score maps (La Joie et al. 2012). Higher w-scores reflect higher voxel-wise atrophy for each patient, adjusted for demographical variables. This approach is particularly useful when performing correlational analyses with smaller samples since findings are adjusted for nuisance covariates during the preprocessing stage, preserving statistical power for the covariate of interest. Voxel-wise frequency mapping of gray matter w-scores across all patients revealed a rostral brain atrophy pattern consistent with the ALS/bvFTD clinicopathological spectrum (**Figure 2A**). Similarly, a standardized scheme was used to assess post-mortem atrophy severity, which was mild overall (**Table 1**), based on coronal brain slabs (Broe et al. 2003).

**Figure 1.**
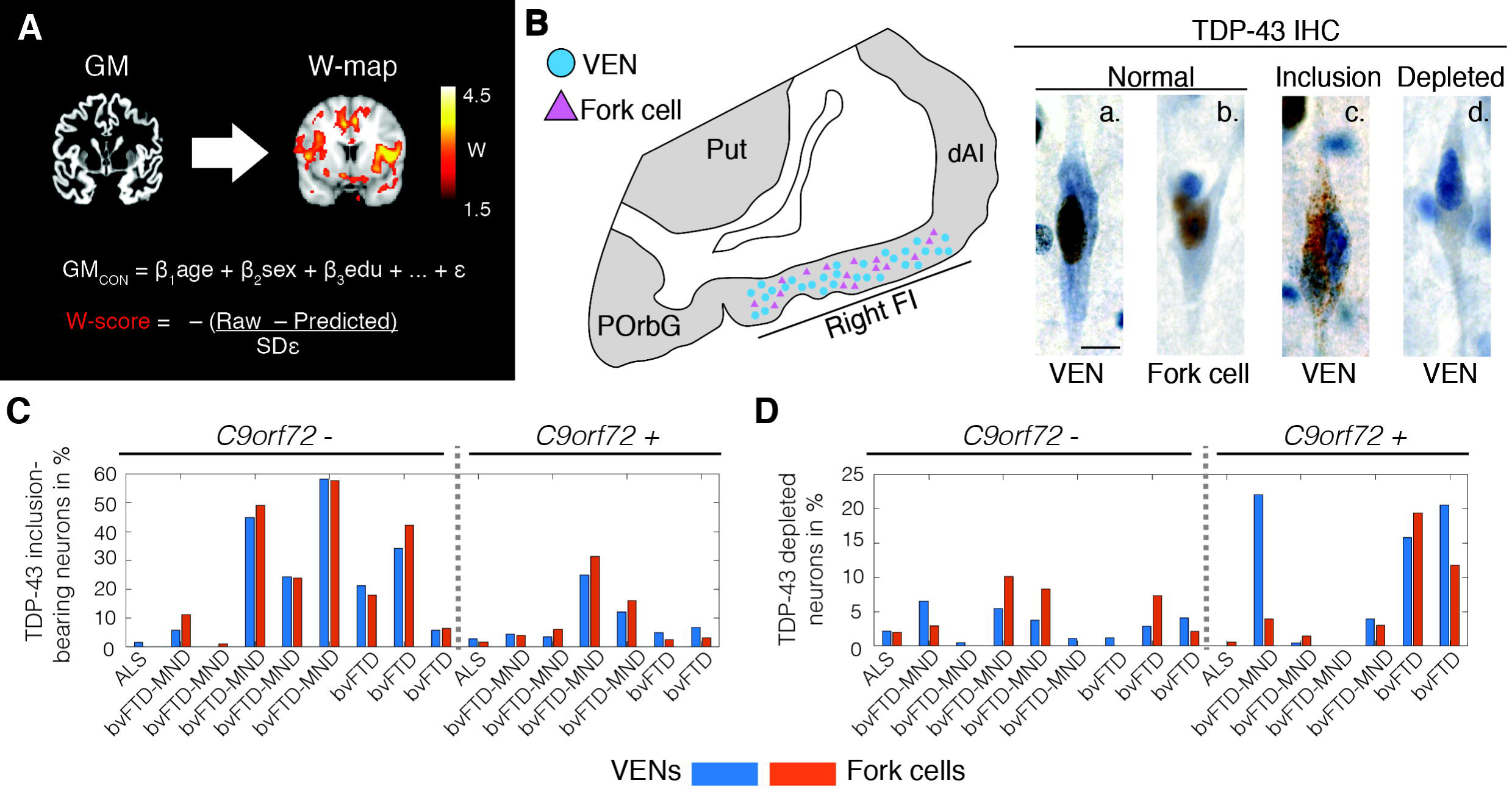
Patient assessment. **(A)** All patients underwent structural MRI. Images were preprocessed, segmented into gray matter tissue intensity maps, and transformed to w-score maps using a multiple regression model trained on healthy older adults (GM_con_) to model the voxel-wise relationship between relevant demographic variables and segmented tissue intensity. The model provides a predicted gray matter map for each patient, and the actual and predicted gray matter maps are compared to derive w-score maps, which reflect the deviation at each voxel from the predicted gray matter intensity. To facilitate the interpretation of these maps, the sign of w-score values was inversed, with higher w-score values reflecting higher levels of atrophy. **(B)** Post-mortem, the rate of VENs (blue circles) and fork cells (violet triangles) with TDP-43 pathobiology was quantified in layer 5 of the right FI via unbiased counting on TDP-43 immunostained sections. TDP-43 is normally expressed in the nucleus (brown nucleus in [a.] normal VEN and [b.] normal fork cell), which in patients aggregates into neuronal cytoplasmic inclusions ([c.] TDP-43 inclusion-bearing VEN) while being cleared from the nucleus. A minority of neurons lacks either normal nuclear TDP-43 or a cytoplasmic inclusion ([d.] nuclear TDP-43 depleted VEN). Scale bar in **B.a**. represents 10 μm. Panel **B** adapted with permission from (Kim et al., 2012). Percentage of **(C)** TDP-43 inclusion-bearing and **(D)** TDP-43 depleted VENs (blue) and fork cells (violet) for individual patients. ALS = amyotrophic lateral sclerosis; bvFTD = behavioral variant frontotemporal dementia; bvFTD-MND = behavioral variant frontotemporal dementia with motor neuron disease; *C9orf72 -* = *C9orf72* expansion non-carriers; *C9orf72* + = *C9orf72* expansion carriers; dAI = dorsal anterior insula; FI = frontoinsular cortex; IHC = immunohistochemistry; GM = gray matter; POrbG = posterior orbitofrontal gyrus; *Predicted* = through the healthy adults regression model predicted patient’s gray matter map; Put = putamen; *Raw* = segmented patient’s gray matter map; *SD*ε = standard deviation of the residuals in the healthy adults model; TDP-43 = transactive response DNA binding protein with 43 kD; VEN = von Economo neuron.

**Figure 2.**
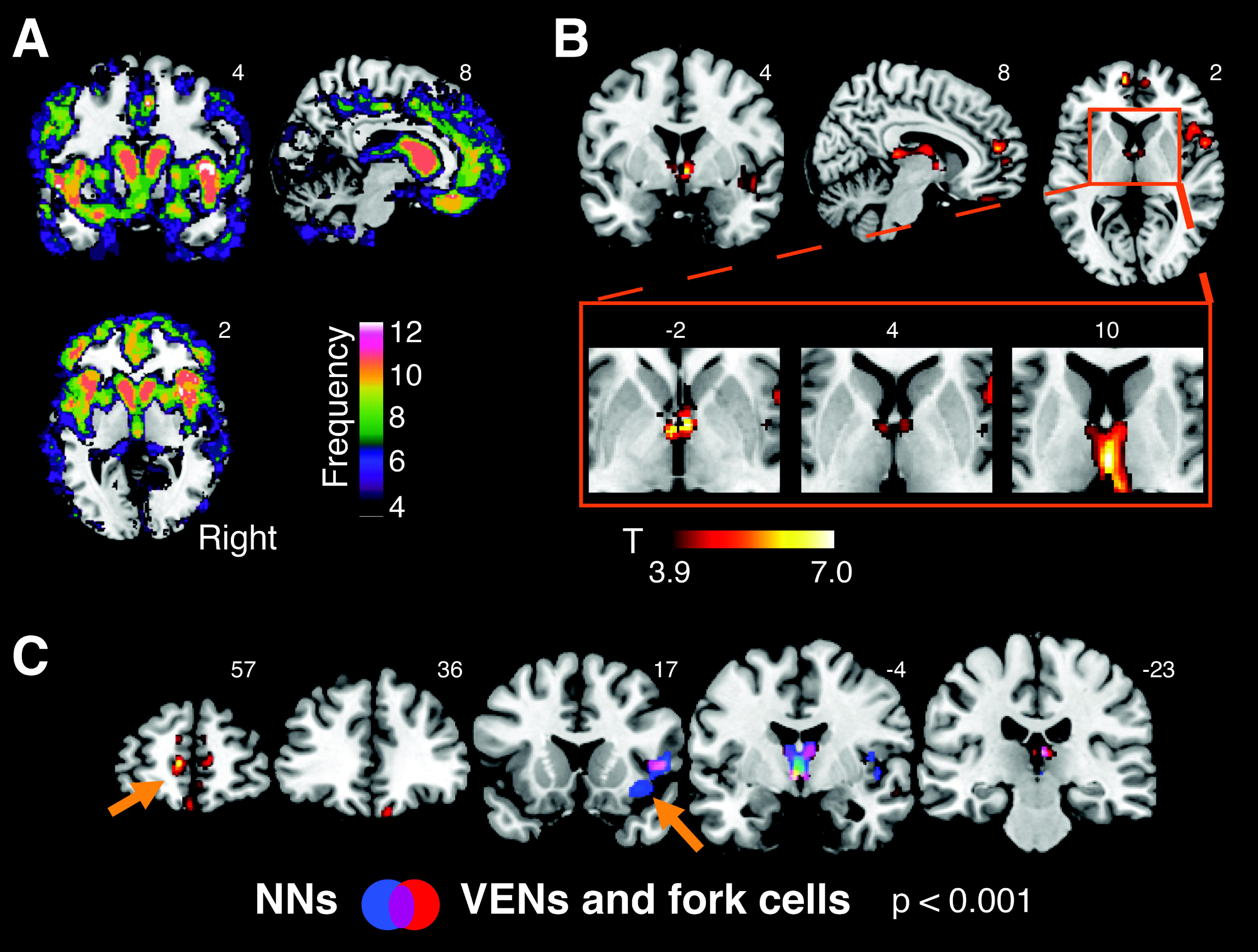
Gray matter atrophy associated with the rate of TDP-43 inclusion-bearing VENs and fork cells in right FI. **(A)** Brain atrophy in ALS/bvFTD spectrum. Voxel-wise frequency map of gray matter atrophy using w-score maps of all patients. Individual w-score gray matter maps were binarized at a w-score threshold of 2 and summed, such that a higher score at each voxel indicates a larger proportion of subjects with suprathreshold atrophy. Only voxels in which at least 4 patients had a w-score > 2 are shown. **(B)** Gray matter atrophy associated with the rate of TDP-43 inclusion-bearing VENs and fork cells in right FI was observed in medial frontal, right insular, subcortical, limbic, and dorsomedial thalamic sites. A cluster in anterior limbic areas is seen in the region of the bed nucleus of the stria terminalis, and subcortical areas span posteriorly into the dorsomedial and medial pulvinar thalamic nuclei. Voxel-wise analyses were corrected for *C9orf72* mutation status and the interval between MRI scan and death. **(C)** Atrophy patterns associated with TDP-43 inclusion formation in VENs and fork cells (in red) versus layer 5 neighboring neurons (in blue) of the right FI. The overlap of both patterns (violet) reveals substantial overlap in right insula and thalamus. Notable differences (orange arrows), include gray matter atrophy in medial frontal regions, including the anterior cingulate cortex, specifically associated with inclusion formation in VENs and fork cells. In contrast, gray matter atrophy of the right FI (where neuron counting was performed) was more associated with inclusion formation in neighboring layer 5 neurons. Voxel-wise analyses were performed with a height threshold of p < 0.001 and a cluster-extent threshold of p < 0.05 FWE, corrected. NNs = layer 5 neighboring neurons.

### VEN and fork cell TDP-43 inclusions in right FI correlate with atrophy in salience network regions

First, we sought to explore the relationship between TDP-43 aggregation in VENs and fork cells, assessed at the cellular level, and brain atrophy, assessed across the whole brain using voxel-based morphometry. Voxel-wise regression analyses were used to identify gray matter structures in which atrophy was related to right FI VEN and fork cell TDP-43 inclusion fraction. Regression models were corrected for *C9orf72* mutation status and scan-to-death interval. This approach revealed that patients with a higher proportion of TDP-43 inclusions in VENs and fork cells showed more severe gray matter atrophy in frontal, insular, limbic and thalamic structures (t = 3.9, height threshold p < 0.001; cluster-extend threshold p < 0.05 FWE corrected); collectively, these structures included key components of the salience network. Specifically, the approach elicited the right FI, as might be expected when using a right FI-derived neuropathological predictor, as well as the pregenual anterior cingulate and paracingulate cortices, medial orbitofrontal cortex, and anterior thalamus, extending into the paraseptal area in the vicinity of the bed nucleus of the stria terminalis and spanning posteriorly the dorsomedial and medial pulvinar thalamic nuclei (**Figure 2B** and **Supplementary Table S2**). Importantly, to assess the influence of using w-score atrophy maps, we performed additional analyses using raw segmented gray matter tissue probability maps (**Supplementary Figure S1** and **Supplementary Table S3**). These analyses yielded findings similar to those produced with the w-score approach, where reduced insular, frontal and subcortical gray matter was associated with higher rates of inclusion formation in VENs and fork cells, as well as a small cluster in the amygdala, another key salience network node.

### Atrophy correlates of VEN and fork cell versus layer 5 neighboring neuron TDP-43 inclusions

The preceding analyses revealed a pattern of gray matter atrophy associated with TDP-43 inclusion-bearing VENs and fork cells. To test the specificity of this association with these neuronal morphotypes, we repeated the preceding analyses using the TDP-43 inclusion fraction in neighboring layer 5 neurons as the predictor. As previously described (Nana et al. 2018), these neighboring neurons showed lower rates of TDP-43 inclusion formation compared to VENs and fork cells (**Table 1)**. For the present sample, the inclusion formation rate for VENs and fork cells was strongly collinear with the rate for neighboring layer 5 neurons (Spearman’s correlation coefficient, Rho = 0.96, p < 0.00001). As in the previous voxel-wise regression analyses, we corrected the model for *C9orf72* mutation status and MRI scan-to-death interval (t = 3.9, height threshold p < 0.001; cluster-extend threshold p < 0.05 FWE corrected). This approach revealed overlapping structural correlates, with subtle but interesting differences noted below. (**Figure 2C** and **Supplementary Table S4**). Similar to VENs and fork cells, the TDP-43 aggregation rate in FI layer 5 neighboring neurons was associated with right insular and medial thalamic gray matter atrophy. In contrast to VENs and fork cells, however, layer 5 neighboring neurons lacked an association with medial frontal/anterior cingulate gray matter atrophy. In the white matter analysis, we uncovered a large cluster in a region abutting the right lateral ventricle but extending into white matter proposed to connect the FI to the anterior cingulate cortex (Cerliani et al. 2012; Ghaziri et al. 2017)(t = 3.9, height threshold p < 0.001; cluster-extent threshold p < 0.05 FWE corrected) (**Supplementary Figure S2** and **Supplementary Table S5**)

### Gray matter atrophy in *C9orf72* expansion carriers is partially accounted by nuclear TDP-43 depletion

In the previous voxel-wise analysis, we identified clusters in which atrophy covaried with VEN and fork cell TDP-43 inclusion formation across all subjects. We were subsequently interested in elucidating how the relationship between atrophy and VEN and fork cell TDP-43 inclusion fraction differed according to *C9orf72* status and clinical syndrome (bvFTD, bvFTD-MND, ALS). We therefore extracted averaged gray matter atrophy levels from clusters associated with VEN and fork cell TDP-43 inclusion fraction and regressed out the influence of interval between MRI scanning and death. To assess the contribution of *C9orf72* expansion carriers to the voxel-wise regression findings, *C9orf72* status was not regressed out from average gray matter atrophy. Partial residuals of gray matter atrophy were then plotted against the rate of TDP-43 inclusion-bearing VENs and fork cells. As expected, patients with pure ALS had fewer VEN and fork cell TDP-43 inclusions compared to patients with bvFTD and bvFTD-MND (**Figure 3A**). Patients with ALS showed the lowest atrophy and fewest inclusion-bearing VENs and fork cells, while patients with bvFTD and bvFTD-MND were spread along the regression line (**Figure 3A**). *C9orf72* expansion carriers had moderate to severe atrophy despite few inclusion-bearing VENs and fork cells compared to expansion non-carriers, suggesting that factors other than TDP-43 aggregation in VENs and fork cells contribute to network-driven regional atrophy in *C9orf72* expansion carriers (Vatsavayai et al. 2016; Nana et al. 2018). Indeed, re-running the preceding voxel-wise regression analysis for the *C9orf72* non-carriers (N = 9) solidified the previous findings and expanded their spatial extent (**Supplementary Figure S3** and **Supplementary Table S7**; t = 5.3, height threshold of p < 0.001 and p < 0.0001; cluster-extent threshold p < 0.05 FWE corrected). Therefore, at least in sporadic bvFTD-ALS spectrum disease, VEN and fork cell TDP-43 inclusion formation may impact an even broader set of brain regions. Why do *C9orf72* mutation carriers show a distinct relationship between gray matter atrophy and TDP-43 pathobiology of VENs and fork cells? Previous work from our group suggests that *C9orf72* expansion carriers, and, to a lesser extent, non-carriers show nuclear TDP-43 depletion, in the absence of cytoplasmic inclusions, and that nuclear depleted cells show equal neuronal atrophy to those bearing an inclusion (Nana et al. 2018). Therefore, to address whether the rate of nuclear TDP-43 depletion influenced the relationship between gray matter atrophy and VEN and fork cell pathobiology, we used a composite measure representing the summed proportion of VENs and fork cells showing (1) inclusion formation with nuclear depletion and (2) nuclear depletion (without inclusion formation) in right FI VENs and fork cells. Voxel-wise analyses on all patients corrected for *C9orf72* status and MRI scan-to-death interval revealed limbic, thalamic, superior parietal, and frontal clusters of gray matter atrophy associated with the VEN and fork cell composite TDP-43 pathobiology score (**Figure 3B** and **Supplementary Table S7**; t = 3.9, height threshold p < 0.001 cluster-extent threshold p < 0.05 FWE corrected). This regional pattern showed substantial overlap with the atrophy pattern associated with the VENs and fork cell TDP-43 inclusion fraction (**Figure 3B)**. Averaged gray matter atrophy was obtained from clusters associated with the TDP-43 composite score, and partial residuals were derived by correcting for MRI scan-to-death interval. These residuals were plotted against the TDP-43 composite score. *C9orf72* mutation carriers were distributed closer to the regression line for the composite score compared to the plot derived using inclusion formation rates alone (**Figure 3C**). These findings suggest that the combination of TDP-43 inclusion formation and nuclear depletion may better account for frontal, limbic, and thalamic atrophy, particularly in *C9orf72* expansion carriers (Nana et al. 2018).

**Figure 3.**
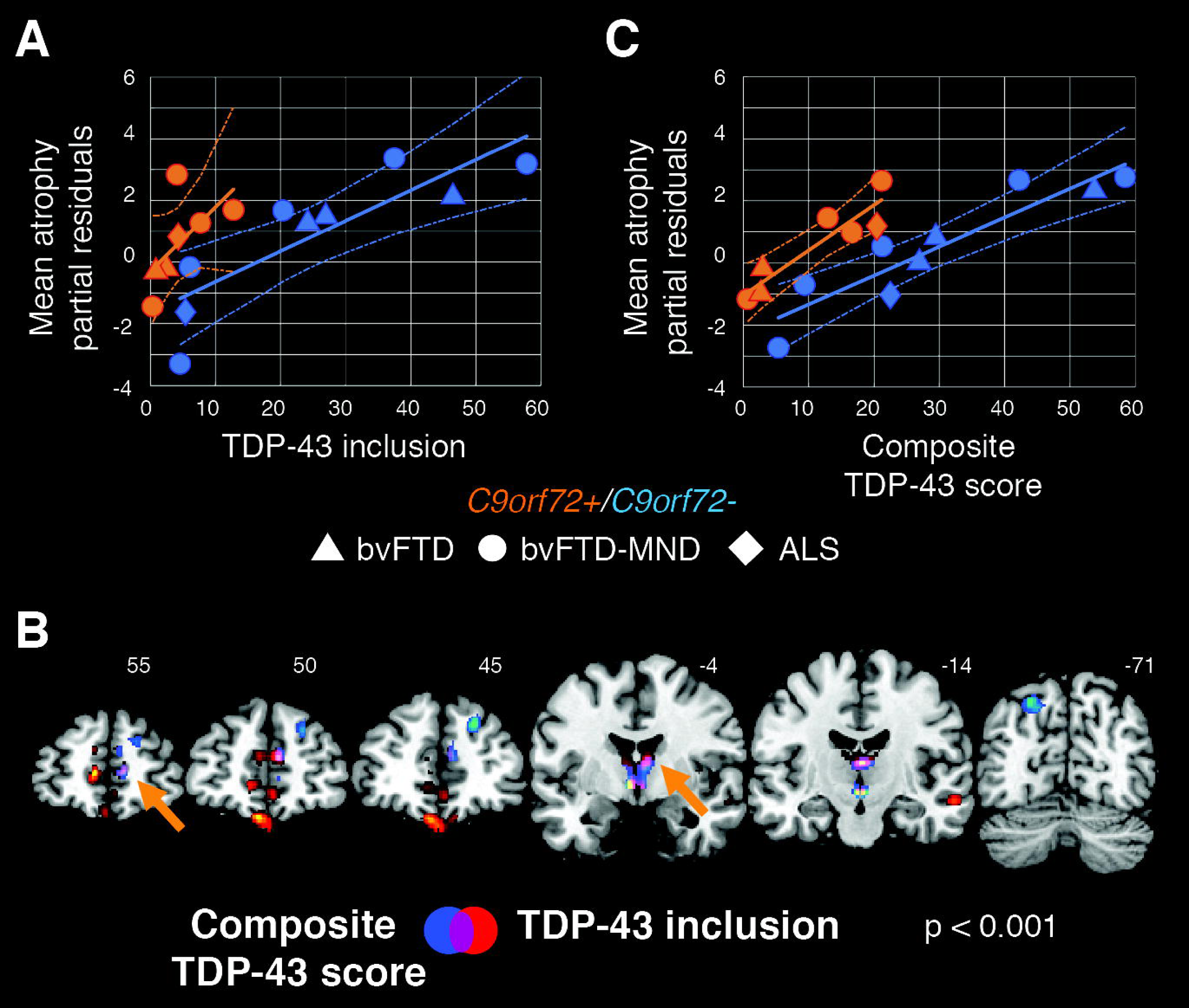
Nuclear TDP-43 depletion partially accounts for gray matter atrophy in *C9orf72* expansion carriers. **(A)** Averaged levels of gray matter atrophy in the identified clusters from **Figure 2B** plotted against the rate of TDP-43 inclusion-bearing VENs and fork cells separately for *C9orf72* expansion carriers and non-carriers. The plotted partial residuals for atrophy were derived by regressing out the influence of interval between scan and death, but not the influence of *C9orf72* status since we were interested in exploring the distribution of *C9orf72* mutation carriers along the regression line. Notably, *C9orf72* expansion carriers (orange) distribute less evenly along the regression line due to high levels of gray matter atrophy despite a low proportion of inclusion-bearing neurons compared to non-expansion carriers (blue). **(B)** Gray matter atrophy in frontal, subcortical limbic, dorsomedial thalamic, and left angular sites is significantly associated with a composite score of VEN/fork cell TDP-43 pathobiology derived by adding the rates of TDP-43 inclusion formation and nuclear depletion (without inclusion) in the right FI VENs and fork cells (in blue). This pattern overlapped substantially (violet and orange arrows) with the atrophy pattern associated with the rate of TDP-43 inclusion-bearing VENs and fork cells (in red, from **Figure 2B**). Voxel-wise analyses were corrected for *C9orf72* mutation status and time interval between MRI scan and death using a height threshold of p < 0.001 and a cluster-extent threshold of p < 0.05 FWE corrected. **(C)** VEN and fork cell TDP-43 composite score plotted against averaged atrophy derived from the clusters identified in panel **B** (blue and violet), after regressing out the influence of interval between scan and death (partial residuals). Combining nuclear TDP-43 depletion with TDP-43 inclusions reveals a more evenly distributed relationship between TDP-43 pathobiology and gray matter atrophy in *C9orf72* expansion carriers and non-carriers than seen when using inclusion-bearing VENs and fork cells alone. Geometric shapes depict distinct FTD syndromes. ALS = amyotrophic lateral sclerosis (diamonds); bvFTD = behavioral variant frontotemporal dementia (triangles); *C9orf72* = chromosome 9 open reading frame 72; bvFTD-MND = frontotemporal dementia with motor neuron disease (circles).

### VEN and fork cell TDP-43 inclusion formation-linked atrophy correlates with deficits in empathic concern

Finally, we sought to relate the atrophy patterns associated with VEN and fork cell TDP-43 inclusion formation with social-emotional deficits, one of the clinical hallmarks of bvFTD (Rankin et al. 2006; Seeley et al. 2012). Social-emotional function was measured using the Interpersonal Reactivity Index (Davis 2007), and we focused on the empathic concern subscale because this measure best reflects the other-centered prosocial response resulting from sharing and understanding another’s emotional state (Rankin et al. 2006), a proposed function of the anterior insula (Lamm and Singer 2010; Leigh et al. 2013). Reflecting the clinical characteristics of each syndrome, in our sample empathic concern was preserved in ALS but severely impaired in bvFTD and bvFTD-MND (**Figure 4A-C**). Spearman partial correlation analyses corrected for *C9orf72* mutation status (**Figure 4A**), revealed that greater deficits in empathic concern were associated with more severe gray matter atrophy averaged across frontal, subcortical, and right insula regions (Rho = -0.86, p < 0.0005). Control analyses were carried out to test for the specificity of these associations. First, we correlated the empathic concern scores with gray matter atrophy derived from a region-of-interest located in the right occipital cortex, and as such beyond the spatial boundaries of the salience network (**Supplementary Figure S4**). Second, we correlated gray matter atrophy averaged across frontal, subcortical, and right insula regions with performance on the Boston Naming Test (Kaplan et al. 1983). These analyses revealed that: *(i)* deficits in empathic concern were not associated with gray matter atrophy in right occipital regions (Rho = -0.19, p = 0.55) (**Figure 4B**); and *(ii)* performance on the Boston Naming Test was not associated with gray matter atrophy averaged across frontal, subcortical, and right insula regions related to VEN and fork cell TDP-43 inclusion fraction (Rho = -0.37, p = 0.24) (**Figure 4C**).

**Figure 4.**
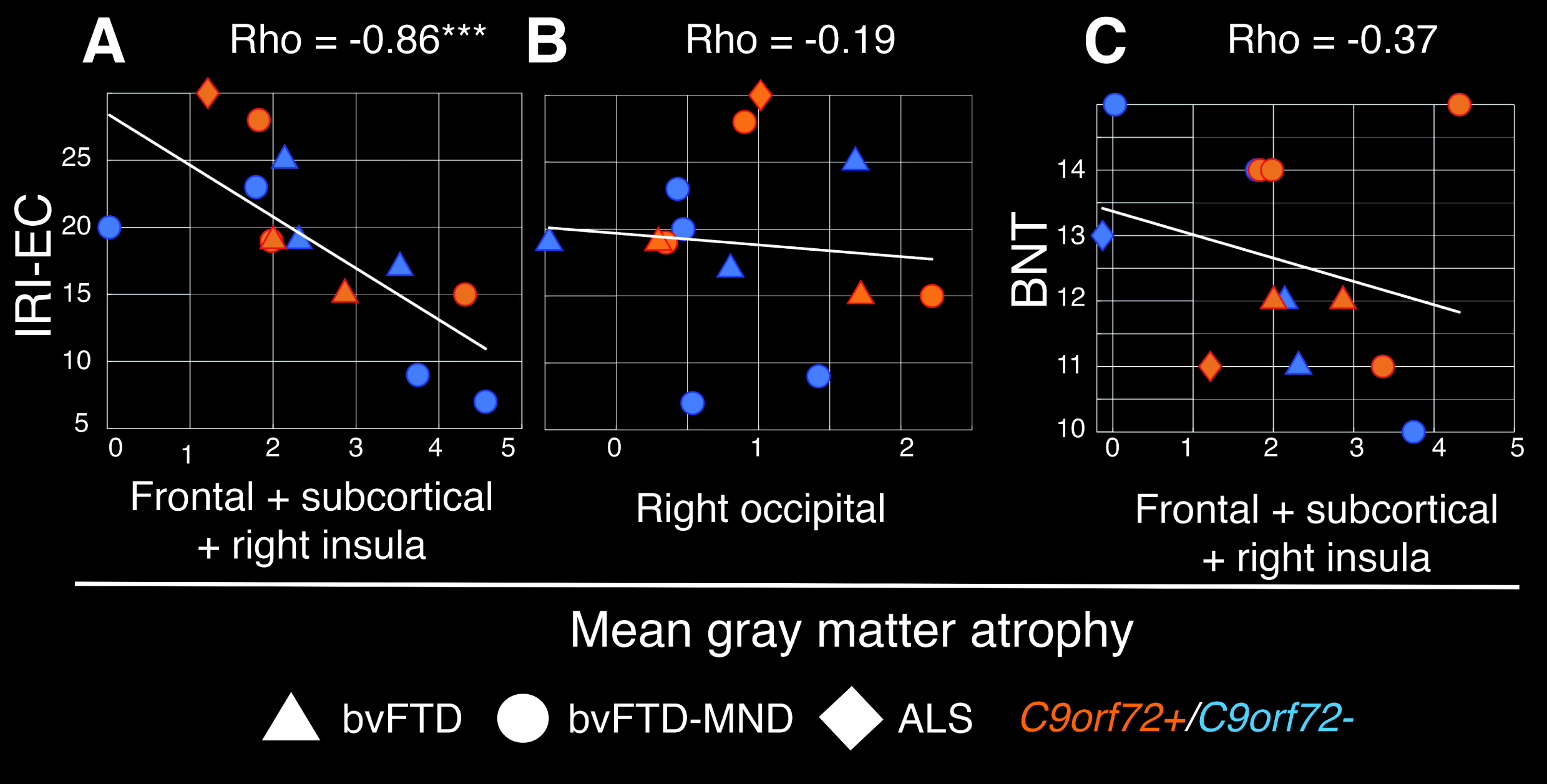
Loss of emotional empathy is associated with atrophy in regions related to VEN and fork cell TDP-43 inclusion formation. **(A)** Gray matter atrophy associated with TDP-43 inclusion-bearing VENs and fork cells (average across clusters identified in **Figure 2B**) correlate with deficits in empathic concern. **(B)** Gray matter atrophy in the right occipital cortex is not significantly associated with deficits in empathic concern. **(C)** Gray matter atrophy associated with TDP-43 inclusion-bearing VENs and fork cells is not significantly associated with performance on the Boston Naming Test. Spearman partial correlation Rho corrected for *C9orf72* mutation status; ***p < 0.0005. BNT = Boston Naming Test; IRI-EC = Interpersonal Reactivity Index, empathic concern subscale.

### Atrophy of salience network regions mediates the link between inclusion-bearing VENs and fork cells and empathy deficits

Empathy deficits are a core feature of bvFTD and have been associated with TDP-43 inclusion formation in VENs and fork cells (Kim et al. 2012; Seeley et al. 2012; Nana et al. 2018) and with structural and functional alterations in brain regions belonging to the salience network (Rankin et al. 2006; Seeley et al. 2012; Toller et al. 2018). In the present study, as in previous work, a linear regression model revealed that VEN and fork cell TDP-43 inclusion formation predicted emotional empathy deficits (**Figure 5A**; c = -0.60, t = -2.47, p < 0.05). We hypothesized that degeneration of salience network regions would mediate the link between neuron type-specific pathobiology and emotional empathy deficits and tested this idea by entering averaged gray matter atrophy derived from the clusters identified in **Figure 2B** in a mediation analysis (Baron and Kenny 1986) (**Figure 5B**). This mediation analysis revealed that TDP-43 inclusion-bearing VENs and fork cells predicted frontal, insular, and thalamic gray matter atrophy averaged across sites as shown in the previous voxel-wise regression analyses (a = 0.61, t = 2.54, p < 0.05). TDP-43 inclusion-bearing VENs and fork cells, however, no longer significantly predicted loss of empathy (c’ = -0.24, t = -0.91, p = 0.38), with average frontal, insular, and thalamic gray matter atrophy fully mediating this relationship (b = -0.59, t = -2.30, p < 0.05).

**Figure 5.**
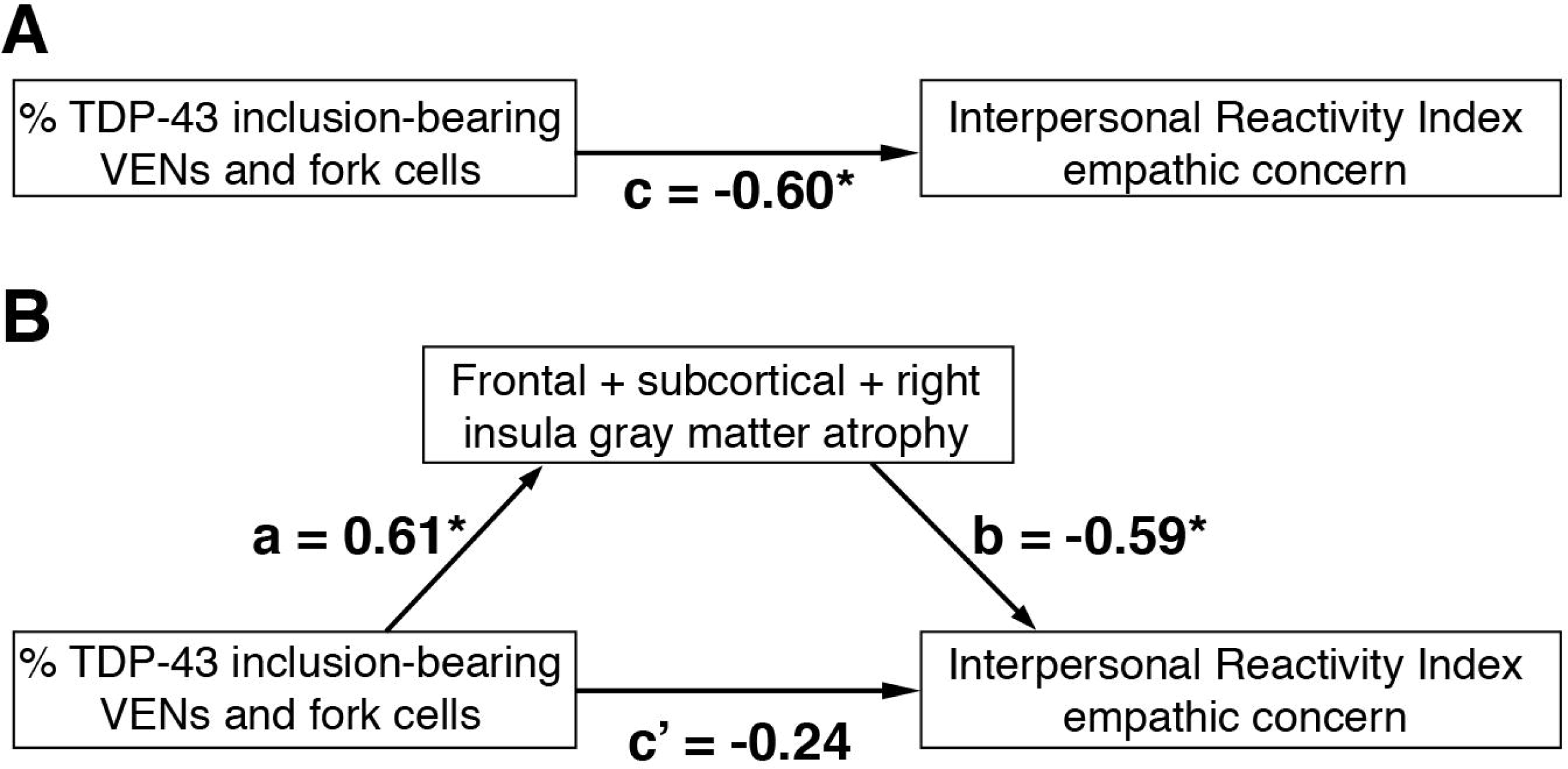
Atrophy of salience network regions mediates the relationships between neuron type-specific TDP-43 pathobiology and loss of emotional empathy. **(A)** As shown in previous work, VEN and fork cell TDP-43 inclusion formation predict empathy loss assessed through the empathic concern subscale of the Interpersonal Reactivity Index. **(B)** The relationship between VEN and fork cell TDP-43 inclusion formation and empathy loss is fully mediated by atrophy of salience network regions, assessed by averaging gray matter w-score values extracted from the clusters identified in **Figure 2B**. a, b, c, and c’ = regression coefficients; *p < 0.05

## Discussion

The ALS/bvFTD clinicopathological continuum linked to TDP-43 proteinopathy provides a unique lesion model for studying the cellular and large-scale network contributors to human social-emotional functioning. Capitalizing on this opportunity, we leveraged a dataset that combined behavioral, neuroimaging, and histopathological data from the same patients with either bvFTD, bvFTD-MND, or ALS. We found that pathobiology of VENs and fork cells, as assessed by the prevalence of TDP-43 inclusions within these neurons, mediated atrophy within key salience network regions, which, in turn, mediated emotional empathy deficits in bvFTD. Patients with bvFTD/ALS due to the *C9orf72* repeat expansion showed a different relationship between TDP-43 inclusion formation, brain atrophy, and empathy, with substantial atrophy despite few TDP-43 inclusions in VENs and fork cells. Interestingly, patients with *C9orf72* repeat expansions may show heightened levels of nuclear TDP-43 depletion without inclusion formation, and adding nuclear TDP-43 depleted VENs and fork cells partially recovered the relationship between VEN/fork cell TDP-43 pathobiology and brain atrophy. Overall, the findings suggest that VENs and fork cells of the frontoinsular cortex play a key role within the salience network (Allman et al. 2011; Seeley et al. 2012), which, in turn, supports emotional empathy (Rankin et al. 2006; Seeley et al. 2012; Toller et al. 2018) likely through domain-general processes related to homeostatic behavioral guidance (Seeley et al. 2012; Critchley and Harrison 2013; Zhou and Seeley 2014; Sturm et al. 2018; Seeley 2019; Pasquini et al. 2020).

### Regional correlates of VEN and fork cell pathobiology: a window into connectivity?

Among commonly studied laboratory mammals, VENs and fork cells are found only in the macaque monkey, and to date little information is available about the connectivity of these large, glutamatergic projection neurons (Allman et al. 2011). Limited tract tracing studies have revealed that VENs in the macaque brain send axons to the contralateral anterior agranular insula and ipsilateral posterior/mid-insula (Evrard et al. 2012). VENs and fork cells express transcription factors associated with subcerebral projections neurons (Cobos and Seeley 2015), suggesting the possibility of projections to autonomic control sites in the brainstem. Given the role of the salience network in viscero-autonomic processing, specific candidate VEN/fork cell projection targets include limbic/subcortical nodes of the salience network, such as the amygdala, hypothalamus, periaqueductal gray matter, and parabrachial nucleus (Cobos and Seeley 2015).

Based on the emerging principles of network-based, transneuronal degeneration, we hypothesize that the structural correlates of VEN/fork cell inclusion formation, as revealed here, provide clues to the likely axonal connections of these cell types, a topic that can be studied only indirectly in humans. At the cortical level, VEN and fork cell inclusion formation was specifically associated with anterior cingulate/paracingulate and medial orbitofrontal degeneration, an association not found for neighboring layer 5 neurons. These regions have structural connections to the right FI (Mesulam 1990), and in humans diffusion-weighted MRI and task-free fMRI studies have revealed structural and functional connections between these regions and FI (Seeley et al. 2007; Cerliani et al. 2012; Uddin 2014; Ghaziri et al. 2017). A direct connection between the FI and the anterior cingulate, mediated by these cell types, would have particular functional significance. The FI has been conceived as the “afferent hub” of the salience network, equipped to build an integrated and contextualized representation of the body, including its responses to prevailing environmental conditions, while the anterior cingulate may serve as the major “efferent hub”, generating those same viscero-autonomic responses to prevailing (internal and external) conditions (Zhou and Seeley 2014). At the subcortical level, VEN and fork cell TDP-43 pathobiology was associated with atrophy in a cluster near the anterior thalamus, possibly extending into the bed nucleus of the stria terminalis. The bed nucleus of the stria terminalis has known projections to salience network nodes such as the centromedial amygdala, orbitofrontal cortex, and anterior cingulate (Lebow and Chen 2016) and has been associated with regulating body homeostasis, stress responses, and social anxiety, facilitating reproductive behavior, and mediating social dysfunction in anxiety disorders and other psychiatric diseases (Lebow and Chen 2016). Mechanisms other than transneural TDP-43 spreading, however, could lead to structural atrophy in regions associated with VEN and fork cell pathobiology, including dysregulated axonal maintenance due to TDP-43 loss of function. For example, recent research has shown that TDP-43 regulates neuronal expression of *STMN2*, a gene involved in axon regeneration and possibly maintenance (Klim et al. 2019). Loss of *STMN2* function due to loss of TDP-43 function could contribute to degeneration of salience network regions by affecting the structural integrity of VENs and fork cell axons.

Three technical limitations need to be considered when interpreting our findings as potential evidence of VEN/fork cell connectivity. First, our study lacked the spatial resolution to delineate some subcortical areas, in particular smaller hypothalamic and brainstem nuclei involved in autonomic processing, so the absence of a relationship to such regions does not provide strong evidence that VENs and fork cells lack connections to those areas. Second, although neighboring layer 5 neurons were not associated with medial frontal and anterior cingulate gray matter atrophy, the overall atrophy maps associated with layer 5 neurons and VENs/fork cells showed substantial overlap. We advise caution in interpreting frontal gray matter atrophy patterns as specific to VEN/fork cell pathobiology, since our assessment of neighboring layer 5 neurons encompassed heterogeneous neuron types, which could have contributed to the differences in structural correlates observed. Third, the specific connection to the anterior cingulate cortex could simply reflect the presence of VENs (and, to a much lesser degree, fork cells) in this region (Gefen et al. 2018); patients showing greater VEN/fork cell involvement in FI may also have had greater involvement of these same vulnerable neurons in the anterior cingulate cortex, independent of spread from FI to this region.

### TDP-43 depletion in VENs and fork cells as a potential atrophy driving mechanism in *C9orf72* expansion carriers

*C9orf72* repeat expansions are the most common genetic cause of FTD and ALS. In bvFTD, *C9orf72* expansion carriers exhibit social-emotional and salience network dysfunction similar to patients with sporadic disease (Boxer et al. 2011; Sha et al. 2012; Irwin et al. 2013). Carriers often show milder atrophy, however, which may be focally accentuated in medial thalamic nuclei (Lee et al. 2014). Patients with *C9orf72*-bvFTD show dramatically fewer VEN and fork cell TDP-43 inclusions despite similar levels of VEN/fork cell dropout (Yang et al. 2017; Nana et al. 2018). On the other hand, carriers often show higher levels of nuclear TDP-43 depletion in the absence of inclusion formation, which is associated with neuronal morphometric changes comparable to that seen in TDP-43 inclusion-bearing neurons (Nana et al. 2018). Our findings show that *C9orf72* expansion carriers have higher than expected frontal, subcortical, and insular atrophy given their low rates of TDP-43 inclusion formation compared to non-carriers. The relationship between TDP-43 pathobiology and regional atrophy was stronger when using a combined measure of TDP-43 inclusion formation and nuclear depletion in VENs and fork cells, suggesting that accounting for all forms of TDP-43 pathobiology better represents the processes leading to atrophy of key salience network hubs.

### Bridging the levels: the neural correlates of social-emotional behavior

Although feature-selective neurons exist within human sensory cortices, complex human experiences and behaviors depend on coordinated activity of distributed brain areas and their underlying neuronal circuit assemblies rather than on firing of single neurons or even neuron types (Mesulam 1990). Previous studies have shown that social-emotional deficits in bvFTD relate (*i*) at the neuronal level, to VEN and fork cells pathobiology (Kim et al. 2012; Nana et al. 2018) and (*ii*) at the large-scale network level, to functional and structural deterioration of the salience network (Rankin et al. 2006; Seeley et al. 2012; Toller et al. 2018). Our findings suggest that pathobiology of specific neuron-types, such as VENs and fork cells, does not directly cause social-emotional deficits but instead leads to degeneration of salience network regions, which, in turn, contributes to loss of emotional empathy, one of the core social-emotional deficits seen in bvFTD. Although little is known about the neuronal mechanisms underlying empathy, neuroimaging studies have associated salience network integrity with social-emotional functioning in both healthy subjects (Shamay-Tsoory 2011; Chételat et al. 2017) and clinical conditions, including sociopathy, Asperger’s syndrome, and schizophrenia (Menon and Uddin 2010). Among salience network nodes, the right FI has been proposed to help coordinate autonomic outflow with highly processed sensory information related to homeostatic, affective, motivational, and hedonic conditions (Craig, 2009; Critchley and Harrison, 2013; Uddin, 2014; Zhou and Seeley, 2014). The anterior and dorsomedial thalamic nuclei have known structural projections to frontal areas such as the anterior cingulate and orbitofrontal cortices (Pergola et al. 2018), two regions that stood out in our findings. Although our results do not inform causal relationships and are derived from a relatively small sample due to the laborious nature of the quantitative neuropathological methods, the novelty of this study relates to the support our data provide for a chain of influence that links specific neuronal morphotypes to brain network structure to emotional empathy, pointing towards the cellular and large-scale brain circuits that are critical for empathy. More research is needed, however, to elucidate how VENs, fork cells, and related neuronal types support the neural circuits underlying diverse aspects of social-emotional function. We hope that the present findings will inform the anatomy of other neuropsychiatric diseases in which loss of emotional empathy is a major feature.

## Abbreviations

ALS: amyotrophic lateral sclerosis;
bv: behavioral variant;
*C9orf72*: chromosome 9 open reading frame 72;
FI: frontoinsular cortex;
FTD: frontotemporal dementia;
MND: motor neuron disease;
MRI: magnetic resonance imaging;
TDP-43: transactive response DNA binding protein 43 kD;
VENs: von Economo neurons

## Acknowledgements

We thank the participants and their families for their invaluable contributions to neurodegeneration research. This study was supported by NIH grants R01AG033017 (WWS), K08 AG052648 (SS), P01AG019724 and P50AG023501 (BLM), and the John Douglas French Alzheimer’s Foundation (LTG). LL was supported by the Reserve Talents of Universities Overseas Research Program of Heilongjiang in China (Document Number: Heijiaogao [2012]381).

## References

Allman JM, Tetreault NA, Hakeem AY, Manaye KF, Semendeferi K, Erwin JM, Park S, Goubert V, Hof PR. 2011. The von Economo neurons in the frontoinsular and anterior cingulate cortex. Ann N Y Acad Sci. 1225:59–71.

Ashburner J, Friston KJ. 2000. Voxel-based morphometry - The methods. Neuroimage. 11:805–821.

Baron RM, Kenny DA. 1986. The Moderator-Mediator Variable Distinction in Social Psychological Research: Conceptual, Strategic, and Statistical Considerations. J Pers Soc Psychol. 51:1173–1182.

Boccardi M, Laakso MP, Bresciani L, Galluzzi S, Geroldi C, Beltramello A, Soininen H, Frisoni GB. 2003. The MRI pattern of frontal and temporal brain atrophy in fronto-temporal dementia. Neurobiol Aging. 24:95–103.

Boxer AL, Mackenzie IR, Boeve BF, Baker M, Seeley WW, Crook R, Feldman H, Hsiung G-YR, Rutherford N, Laluz V, Whitwell J, Foti D, McDade E, Molano J, Karydas A, Wojtas A, Goldman J, Mirsky J, Sengdy P, DeArmond S, Miller BL, Rademakers R. 2011. Clinical, neuroimaging and neuropathological features of a new chromosome 9p-linked FTD-ALS family. J Neurol Neurosurg Psychiatry. 82:196–203.

Brettschneider J, Del Tredici K, Lee VM-Y, Trojanowski JQ. 2015. Spreading of pathology in neurodegenerative diseases: a focus on human studies. Nat Rev Neurosci. 16:109–120.

Broe M, Hodges JR, Schofield E, Shepherd CE, Kril JJ, Halliday GM. 2003. Staging disease severity in pathologically confirmed cases of frontotemporal dementia. Neurology. 60:1005–1011.

Brown JA, Neuhaus J, Miller BL, Rosen HJ, Seeley WW, Brown JA, Neuhaus J, Sible IJ, Sias AC, Lee SE, Kornak J, Marx GA, Karydas AM, Spina S, Grinberg LT, Coppola G, Geschwind DH, Kramer JH, Gorno-tempini ML, Miller BL, Rosen HJ, Seeley WW. 2019. Patient-Tailored, Connectivity-Based Forecasts of Spreading Brain Atrophy. Neuron. 1–13.

Cerliani L, Thomas RM, Jbabdi S, Siero JCW, Nanetti L, Crippa A, Gazzola V, D’Arceuil H, Keysers C. 2012. Probabilistic tractography recovers a rostrocaudal trajectory of connectivity variability in the human insular cortex. Hum Brain Mapp. 33:2005–2034.

Chételat G, Mézenge F, Tomadesso C, Landeau B, Arenaza-urquijo E, Rauchs G, André C, Flores R De, Egret S, Gonneaud J, Poisnel G, Chocat A, Quillard A, Desgranges B, Bloch J, Ricard M, Lutz A. 2017. Reduced age-associated brain changes in expert meditators□: a multimodal neuroimaging pilot study. Sci Rep. 1–11.

Cobos I, Seeley WW. 2015. Human von economo neurons express transcription factors associated with Layer v subcerebral projection neurons. Cereb Cortex. 25:213–220.

Cope TE, Shtyrov Y, MacGregor LJ, Holland R, Pulvermüller F, Rowe JB, Patterson K. 2020. Anterior temporal lobe is necessary for efficient lateralised processing of spoken word identity. Cortex. 126:107–118.

Craig AD. 2009. How do you feel — now? The anterior insula and human awareness. Nat Rev Neurosci. 10:59–70.

Critchley HD, Harrison NA. 2013. Visceral Influences on Brain and Behavior. Neuron. 77:624–638.

Davis MHA. 2007. A Multidimensional Approach to Individual Differences in Empathy. Journal of Personality and Social Psychology

DeJesus-Hernandez M, Mackenzie IR, Boeve BF, Boxer AL, Baker M, Rutherford NJ, Nicholson AM, Finch NA, Flynn H, Adamson J, Kouri N, Wojtas A, Sengdy P, Hsiung GYR, Karydas A, Seeley WW, Josephs KA, Coppola G, Geschwind DH, Wszolek ZK, Feldman H, Knopman DS, Petersen RC, Miller BL, Dickson DW, Boylan KB, Graff-Radford NR, Rademakers R. 2011. Expanded GGGGCC Hexanucleotide Repeat in Noncoding Region of C9ORF72 Causes Chromosome 9p-Linked FTD and ALS. Neuron. 72:245–256.

Evrard HC, Forro T, Logothetis NK. 2012. Von Economo Neurons in the Anterior Insula of the Macaque Monkey. Neuron. 74:482–489.

Fan L, Li H, Zhuo J, Zhang Y, Wang J, Chen L, Yang Z, Chu C, Xie S, Laird AR, Fox PT, Eickhoff SB, Yu C, Jiang T. 2016. The Human Brainnetome Atlas: A New Brain Atlas Based on Connectional Architecture. Cereb Cortex. 26:3508–3526.

Gefen T, Papastefan ST, Rezvanian A, Bigio EH, Weintraub S, Rogalski E, Mesulam MM, Geula C. 2018. Von Economo neurons of the anterior cingulate across the lifespan and in Alzheimer’s disease. Cortex. 99:69–77.

Ghaziri J, Tucholka A, Girard G, Houde J-C, Boucher O, Gilbert G, Descoteaux M, Lippé S, Rainville P, Nguyen DK. 2017. The Corticocortical Structural Connectivity of the Human Insula. Cereb Cortex. 27:1216–1228.

Ghosh BCP, Calder AJ, Peers P V., Lawrence AD, Acosta-Cabronero J, Pereira JM, Hodges JR, Rowe JB. 2012. Social cognitive deficits and their neural correlates in progressive supranuclear palsy. Brain. 135:2089–2102.

Gorgolewski KJ, Varoquaux G, Rivera G, Schwarz Y, Ghosh SS, Maumet C, Sochat V V., Nichols TE, Poldrack RA, Poline JB, Yarkoni T, Margulies DS. 2015. NeuroVault.Org: A web-based repository for collecting and sharing unthresholded statistical maps of the human brain. Front Neuroinform. 9:1–9.

Hughes LE, Rittman T, Robbins TW, Rowe JB. 2018. Reorganization of cortical oscillatory dynamics underlying disinhibition in frontotemporal dementia. Brain. 141:2486–2499.

Irwin DJ, McMillan CT, Brettschneider J, Libon DJ, Powers J, Rascovsky K, Toledo JB, Boller A, Bekisz J, Chandrasekaran K, Wood EM, Shaw LM, Woo JH, Cook PA, Wolk DA, Arnold SE, Van Deerlin VM, McCluskey LF, Elman L, Lee VM-Y, Trojanowski JQ, Grossman M. 2013. Cognitive decline and reduced survival in *C9orf72* expansion frontotemporal degeneration and amyotrophic lateral sclerosis. J Neurol Neurosurg Psychiatry. 84:163–169.

Jack CR, Petersen RC, Xu YC, Waring SC, O’Brien PC, Tangalos EG, Smith GE, Ivnik RJ, Kokmen E, Kokmen E. 1997. Medial temporal atrophy on MRI in normal aging and very mild Alzheimer’s disease. Neurology. 49:786–794.

Kaplan E, Goodglass H, Weintraub S. 1983. The Boston Naming Test. 2nd Editio. ed.

Kim EJ, Sidhu M, Gaus SE, Huang EJ, Hof PR, Miller BL, DeArmond SJ, Seeley WW. 2012. Selective frontoinsular von economo neuron and fork cell loss in early behavioral variant frontotemporal dementia. Cereb Cortex. 22:251–259.

Klim JR, Williams LA, Limone F, Guerra San Juan I, Davis-Dusenbery BN, Mordes DA, Burberry A, Steinbaugh MJ, Gamage KK, Kirchner R, Moccia R, Cassel SH, Chen K, Wainger BJ, Woolf CJ, Eggan K. 2019. ALS-implicated protein TDP-43 sustains levels of STMN2, a mediator of motor neuron growth and repair. Nat Neurosci. 22:167–179.

Knopman DS, Kramer JH, Boeve BF, Caselli RJ, Graff-Radford NR, Mendez MF, Miller BL, Mercaldo N. 2008. Development of methodology for conducting clinical trials in frontotemporal lobar degeneration. Brain. 131:2957–2968.

La Joie R, Perrotin A, Barré L, Hommet C, Mezenge F, Ibazizene M, Camus V, Abbas A, Landeau B, Guilloteau D, de La Sayette V, Eustache F, Desgranges B, Chételat G. 2012. Region-Specific Hierarchy between Atrophy, Hypometabolism, and -Amyloid (A) Load in Alzheimer’s Disease Dementia. J Neurosci. 32:16265–16273.

Lamm C, Singer T. 2010. The role of anterior insular cortex in social emotions. Brain Struct Funct. 214:579–591.

Lebow MA, Chen A. 2016. Overshadowed by the amygdala□: the bed nucleus of the stria terminalis emerges as key to psychiatric disorders. Mol Psychiatry. 21:450–463.

Lee SE, Khazenzon AM, Trujillo AJ, Guo CC, Yokoyama JS, Sha SJ, Takada LT, Karydas AM, Block NR, Coppola G, Pribadi M, Geschwind DH, Rademakers R, Fong JC, Weiner MW, Boxer AL, Kramer JH, Rosen HJ, Miller BL, Seeley WW. 2014. Altered network connectivity in frontotemporal dementia with C9orf72 hexanucleotide repeat expansion. Brain. 137:3047–3060.

Leigh R, Oishi K, Hsu J, Lindquist M, Gottesman RF, Jarso S, Crainiceanu C, Mori S, Hillis AE. 2013. Acute lesions that impair affective empathy. Brain. 136:2539–2549.

Mackenzie IRA, Neumann M. 2016. Molecular neuropathology of frontotemporal dementia: insights into disease mechanisms from postmortem studies. J Neurochem. 138:54–70.

MacKenzie IRA, Neumann M, Bigio EH, Cairns NJ, Alafuzoff I, Kril J, Kovacs GG, Ghetti B, Halliday G, Holm IE, Ince PG, Kamphorst W, Revesz T, Rozemuller AJM, Kumar-Singh S, Akiyama H, Baborie A, Spina S, Dickson DW, Trojanowski JQ, Mann DMA. 2010. Nomenclature and nosology for neuropathologic subtypes of frontotemporal lobar degeneration: An update. Acta Neuropathol. 119:1–4.

McKeith IG, Boeve BF, Dickson DW, Halliday G, Taylor J-P, Weintraub D, Aarsland D, Galvin J, Attems J, Ballard CG, Bayston A, Beach TG, Blanc F, Bohnen N, Bonanni L, Bras J, Brundin P, Burn D, Chen-Plotkin A, Duda JE, El-Agnaf O, Feldman H, Ferman TJ, ffytche D, Fujishiro H, Galasko D, Goldman JG, Gomperts SN, Graff-Radford NR, Honig LS, Iranzo A, Kantarci K, Kaufer D, Kukull W, Lee VMY, Leverenz JB, Lewis S, Lippa C, Lunde A, Masellis M, Masliah E, McLean P, Mollenhauer B, Montine TJ, Moreno E, Mori E, Murray M, O’Brien JT, Orimo S, Postuma RB, Ramaswamy S, Ross OA, Salmon DP, Singleton A, Taylor A, Thomas A, Tiraboschi P, Toledo JB, Trojanowski JQ, Tsuang D, Walker Z, Yamada M, Kosaka K. 2017. Diagnosis and management of dementia with Lewy bodies. Neurology. 89:88–100.

McKeith IG, Dickson DW, Lowe J, Emre M, O’Brien JT, Feldman H, Cummings J, Duda JE, Lippa C, Perry EK, Aarsland D, Arai H, Ballard CG, Boeve B, Burn DJ, Costa D, Del Ser T, Dubois B, Galasko D, Gauthier S, Goetz CG, Gomez-Tortosa E, Halliday G, Hansen LA, Hardy J, Iwatsubo T, Kalaria RN, Kaufer D, Kenny RA, Korczyn A, Kosaka K, Lee VMY, Lees A, Litvan I, Londos E, Lopez OL, Minoshima S, Mizuno Y, Molina JA, Mukaetova-Ladinska EB, Pasquier F, Perry RH, Schulz JB, Trojanowski JQ, Yamada M, Consortium on DLB. 2005. Diagnosis and management of dementia with Lewy bodies: Third report of the DLB consortium. Neurology. 65:1863–1872.

Menon V, Uddin LQ. 2010. Saliency, switching, attention and control: a network model of insula function. Brain Struct Funct. 1–13.

Mesulam M-M. 1990. Large-scale neurocognitive networks and distributed processing for attention, language, and memory. Ann Neurol. 28:597–613.

Montine TJ, Phelps CH, Beach TG, Bigio EH, Cairns NJ, Dickson DW, Duyckaerts C, Frosch MP, Masliah E, Mirra SS, Nelson PT, Schneider JA, Thal DR, Trojanowski JQ, Vinters H V, Hyman BT, National Institute on Aging, Alzheimer’s Association. 2012. National Institute on Aging-Alzheimer’s Association guidelines for the neuropathologic assessment of Alzheimer’s disease: a practical approach. Acta Neuropathol. 123:1–11.

Nana AL, Sidhu M, Gaus SE, Hwang J-HL, Li L, Park Y, Kim E-J, Pasquini L, Allen IE, Rankin KP, Toller G, Kramer JH, Geschwind DH, Coppola G, Huang EJ, Grinberg LT, Miller BL, Seeley WW. 2018. Neurons selectively targeted in frontotemporal dementia reveal early stage TDP-43 pathobiology. Acta Neuropathol. 137:27–46.

Neumann M, Sampathu DM, Kwong LK, Truax AC, Micsenyi MC, Chou TT, Bruce J, Schuck T, Grossman M, Clark CM, McCluskey LF, Miller BL, Masliah E, Mackenzie IR, Feldman H, Feiden W, Kretzschmar HA, Trojanowski JQ, Lee VM-Y. 2006. Ubiquitinated TDP-43 in Frontotemporal Lobar Degeneration and Amyotrophic Lateral Sclerosis. Science (80-). 314:130–133.

Nimchinsky EA, Vogt BA, Morrison JH, Hof PR. 1995. Spindle neurons of the human anterior cingul. Ate cortex. J Comp Neurol. 355:27–37.

Pasquini L, Palhano-Fontes F, Araujo DB. 2020. Subacute effects of the psychedelic ayahuasca on the salience and default mode networks. medRxiv. https://doi.org/10.1101/19007542.

Pasquini L, Scherr M, Tahmasian M, Meng C, Myers NE, Ortner M, Mühlau M, Kurz A, Förstl H, Zimmer C, Grimmer T, Wohlschläger AM, Riedl V, Sorg C. 2015. Link between hippocampus’ raised local and eased global intrinsic connectivity in AD. Alzheimer’s Dement. 11:475–84.

Pasquini L, Toller G, Staffaroni A, Brown JA, Deng J, Lee A, Kurcyus K, Shdo SM, Allen I, Sturm VE, Cobigo Y, Borghesani V, Battistella G, Gorno-Tempini ML, Rankin KP, Kramer J, Rosen HH, Miller BL, Seeley WW. 2019. State and trait characteristics of anterior insula time-varying functional connectivity. Neuroimage. 208:716720.

Pergola G, Danet L, Pitel AL, Carlesimo GA, Segobin S, Pariente J, Suchan B, Mitchell AS, Barbeau EJ. 2018. The Regulatory Role of the Human Mediodorsal Thalamus. Trends Cogn Sci. 22:1011–1025.

Raj A, Kuceyeski A, Weiner M. 2012. A Network Diffusion Model of Disease Progression in Dementia. Neuron. 73:1204–1215.

Rankin KP, Gorno-Tempini ML, Allison SC, Stanley CM, Glenn S, Weiner MW, Miller BL. 2006. Structural anatomy of empathy in neurodegenerative disease. Brain. 129:2945–2956.

Renton AE, Majounie E, Waite A, Simón-Sánchez J, Rollinson S, Gibbs JR, Schymick JC, Laaksovirta H, van Swieten JC, Myllykangas L, Kalimo H, Paetau A, Abramzon Y, Remes AM, Kaganovich A, Scholz SW, Duckworth J, Ding J, Harmer DW, Hernandez DG, Johnson JO, Mok K, Ryten M, Trabzuni D, Guerreiro RJ, Orrell RW, Neal J, Murray A, Pearson J, Jansen IE, Sondervan D, Seelaar H, Blake D, Young K, Halliwell N, Callister JB, Toulson G, Richardson A, Gerhard A, Snowden J, Mann D, Neary D, Nalls MA, Peuralinna T, Jansson L, Isoviita V-M, Kaivorinne A-L, Hölttä-Vuori M, Ikonen E, Sulkava R, Benatar M, Wuu J, Chiò A, Restagno G, Borghero G, Sabatelli M, Heckerman D, Rogaeva E, Zinman L, Rothstein JD, Sendtner M, Drepper C, Eichler EE, Alkan C, Abdullaev Z, Pack SD, Dutra A, Pak E, Hardy J, Singleton A, Williams NM, Heutink P, Pickering-Brown S, Morris HR, Tienari PJ, Traynor BJ, Traynor BJ. 2011. A Hexanucleotide Repeat Expansion in C9ORF72 Is the Cause of Chromosome 9p21-Linked ALS-FTD. Neuron. 72:257–268.

Santillo AF, Nilsson C, Englund E. 2013. von Economo neurones are selectively targeted in frontotemporal dementia. Neuropathol Appl Neurobiol. 39:572–579.

Seeley W. 2017. Mapping Neurodegenerative Disease Onset and Progression. Cold Spring Harb Perspect Biol. 1389.

Seeley WW. 2019. The salience network□: a neural system for perceiving and responding to homeostatic demands. J Neurosci. 39:9878–9882.

Seeley WW, Menon V, Schatzberg AF, Keller J, Glover GH, Kenna H, Reiss AL, Greicius MD. 2007. Dissociable intrinsic connectivity networks for salience processing and executive control. J Neurosci. 27:2349–2356.

Seeley WW, Zhou J, Kim E-J. 2012. Frontotemporal Dementia: What Can the Behavioral Variant Teach Us about Human Brain Organization? Neurosci. 18:373–385.

Sha SJ, Takada LT, Rankin KP, Yokoyama JS, Rutherford NJ, Fong JC, Khan B, Karydas A, Baker MC, DeJesus-Hernandez M, Pribadi M, Coppola G, Geschwind DH, Rademakers R, Lee SE, Seeley W, Miller BL, Boxer AL. 2012. Frontotemporal dementia due to C9ORF72 mutations: Clinical and imaging features. Neurology. 79:1002–1011.

Shamay-Tsoory SG. 2011. The Neural Bases for Empathy. Neurosci. 17:18–24.

Silver M, Montana G, Nichols TE. 2011. False positives in neuroimaging genetics using voxel-based morphometry data. Neuroimage. 54:992–1000.

Stimpson CD, Tetreault NA, Allman JM, Jacobs B, Butti C, Hof PR, Sherwood CC. 2011. Biochemical specificity of von economo neurons in hominoids. Am J Hum Biol. 23:22–28.

Sturm VE, Brown JA, Hua AY, Lwi SJ, Zhou J, Kurth F, Eickhoff SB, Rosen HJ, Kramer JH, Miller BL, Levenson RW, Seeley WW. 2018. Network architecture underlying basal autonomic outflow: Evidence from frontotemporal dementia. J Neurosci. 0347–18.

Tartaglia MC, Sidhu M, Laluz V, Racine C, Rabinovici GD, Creighton K, Karydas A, Rademakers R, Huang EJ, Miller BL, DeArmond SJ, Seeley WW. 2010. Sporadic corticobasal syndrome due to FTLD-TDP. Acta Neuropathol. 119:365–374.

Toller G, Brown J, Sollberger M, Shdo S, Bouvet L, Sukhanov P, Seeley WW, Miller BL, Rankin KP. 2018. Individual differences in socioemotional sensitivity are an index of salience network function. Cortex. 1–13.

Uddin LQ. 2014. Salience processing and insular cortical function and dysfunction. Nat Rev Neurosci. 16:55–61.

Vatsavayai SC, Yoon SJ, Gardner RC, Gendron TF, Vargas JNS, Trujillo A, Pribadi M, Phillips JJ, Gaus SE, Hixson JD, Garcia PA, Rabinovici GD, Coppola G, Geschwind DH, Petrucelli L, Miller BL, Seeley WW. 2016. Timing and significance of pathological features in C9orf72 expansion-associated frontotemporal dementia. 139:3202–3216.

Yang Y, Halliday GM, Hodges JR, Tan RH. 2017. Von Economo Neuron Density and Thalamus Volumes in Behavioral Deficits in Frontotemporal Dementia Cases with and without a C9ORF72 Repeat Expansion. J Alzheimer’s Dis. 58:701–709.

Zhou J, Gennatas ED, Kramer JH, Miller BL, Seeley WW. 2012. Predicting Regional Neurodegeneration from the Healthy Brain Functional Connectome. Neuron. 73:1216–1227.

Zhou J, Seeley WW. 2014. Network Dysfunction in Alzheimer’s Disease and Frontotemporal Dementia□: Implications for Psychiatry. Biol Psychiatry. 75:565–573.

