## supplement for "Salience network atrophy links neuron type-specific pathobiology to loss of empathy in frontotemporal dementia"

**Supplement Information**

**Supplementary Data**

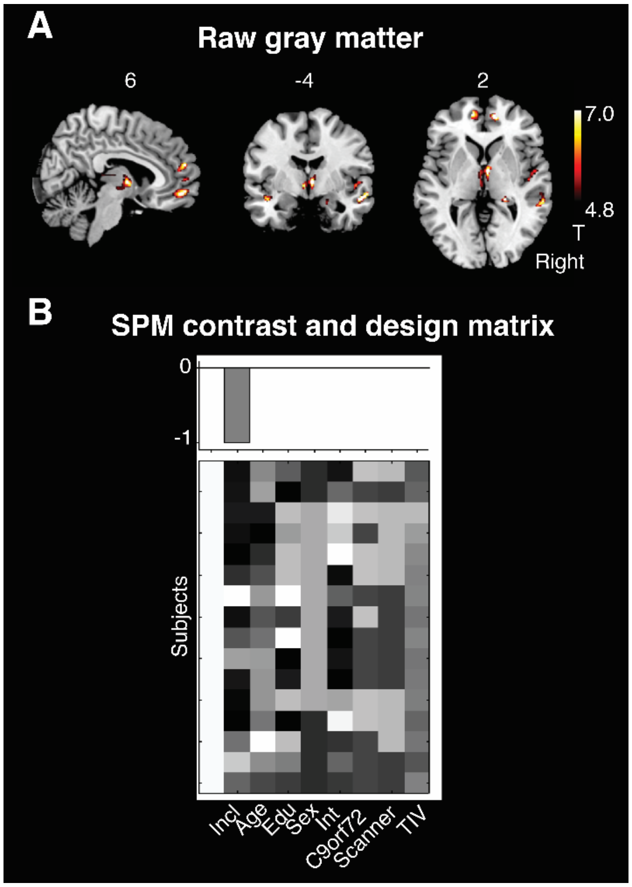


**Figure S1. Regression analyses on unadjusted tissue intensity maps. (A)** Voxel-wise regression analyses on unadjusted, segmented gray matter tissue intensity maps, using the rate of TDP-43 inclusion-bearing VENs and fork cells as regressor ("Incl" term in the contrast model). **(B)** Analyses were corrected for the same variables implemented in the generation of w-score maps: age at scanning (Age, in years), education (Edu, in years), sex, scan-to-death interval (Int, in years), *C9orf72* mutation status, scanner type, and total intracranial volume (TIV, in liters). A covariate for handedness was not added, since all patients participating in the study were right handed. Voxel-wise analyses were performed with a height threshold of $p < 0.001$ and a cluster-extent threshold of $p < 0.05$ FWE corrected.

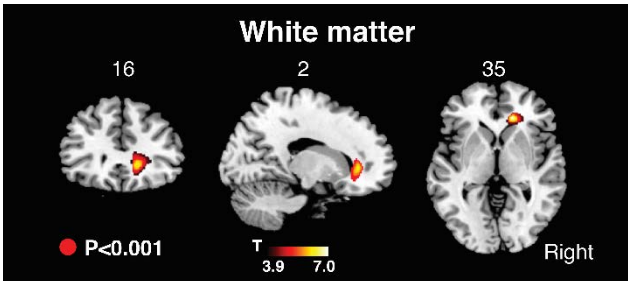


**Figure S2**. **White matter atrophy associated with the rate of TDP-43 inclusion-bearing VENs and fork cells in right FI.** Associated white matter atrophy consisted of a cluster in proximity to the anterior cingulate cortex and right anterior insula.

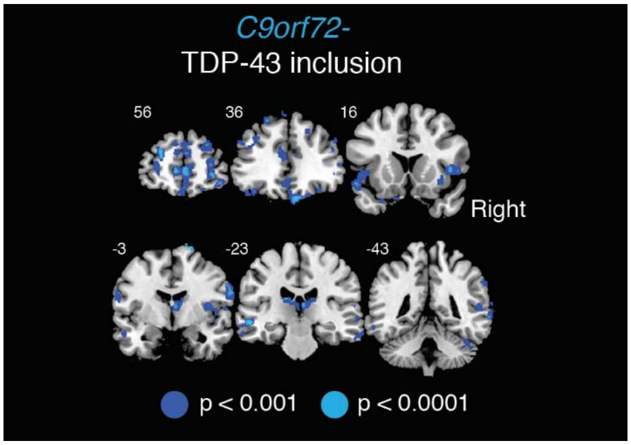


**Figure S3**. **Control voxel-wise analyses without *C9orf72* expansion carriers**. Voxel-wise analyses without *C9orf72* expansion carriers revealed more widespread patterns of gray matter atrophy associated with TDP-43 inclusion-bearing VENs and fork cells. Height thresholds of $p < 0.001$ and $p < 0.0001$, cluster-extent threshold of $p < 0.05$ FWE corrected.

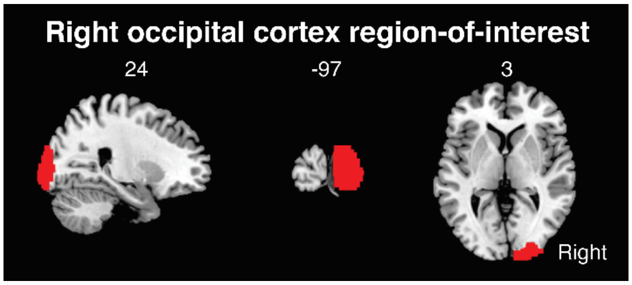


**Figure S4. Region-of-interest in the right occipital cortex**. Averaged levels of gray matter atrophy were derived from this region-of-interest and associated with the empathic concern score of the Interpersonal Reactivity Index.

**Table S1.** Healthy control sample used to build the w-score model. Related to **Methods and Supplementary Experimental Procedures.**

| | |
|---|---|
| N | 288 |
| Age in years | 66.3 (10.8) |
| Gender (Female:Male) | 172:116 |
| Scanner type (1.5T:3T) | 68:220 |
| Handedness (Left:Ambidextrous:Right) | 35:3:250 |
| Education in years | 17.0 (2.0) |
| Total intracranial volume in liters | 1.4 (0.1) |

Unless otherwise noted, numbers in parentheses are SD of preceding mean

**Table S2.** MNI coordinates for cluster peaks in **Figures 2A**.

| Brain Regions | Cluster size (voxels) | MNI peak voxel X Y Z | t-values peak voxel |
|---|---|---|---|
| **Gray matter atrophy associated with inclusion-bearing VENs and fork cells (Figure 2A)** | | | |
| Left ACC | 2060 | -8 57 4 | 6.7 |
| Left OFC | | -6 49 -24 | 6.7 |
| Right ACC | | 8 48 14 | 6.1 |
| Right Thalamus | 1662 | 2 -18 10 | 6.7 |
| Right Thalamus | | 3 0 -2 | 6.7 |
| Right Thalamus | | 2 -10 -9 | 6.7 |
| Right Insula | 1277 | 50 10 -4 | 5.7 |
| Right RO | | 56 8 4 | 5.7 |
| Right MTG | | 58 -12 -14 | 5.7 |

Brain regions were identified on the Harvard-Oxford Cortical and Subcortical Structural Atlases using the peak-voxel MNI coordinates. ACC = anterior cingulate cortex; MTG = medial temporal gyrus; OFC = orbitofrontal cortex; RO = rolandic operculum.

**Table S3.** MNI coordinates for cluster peaks in **Supplementary Figures S1A.**

| Brain Regions | Cluster size (voxels) | MNI peak voxel X Y Z | t-values peak voxel |
|---|---|---|---|
| **Raw gray matter atrophy associated with inclusion-bearing VENs and fork cells *** | | | |
| Right STG | 781 | 60 -3 -10 | 11.5 |
| Right Insula | | 54 -6 -15 | 10.0 |
| Right STG | | 60 -24 -3 | 8.9 |
| Unclassified | 601 | -2 -14 -8 | 11.2 |
| Left Thalamus | | -2 -21 -3 | 9.8 |
| Right Thalamus | | 4 -4 2 | 9.5 |
| Right ACC | 561 | 12 51 2 | 11.1 |
| Right ACC | | 10 52 15 | 10.9 |
| Right MPG | | 8 52 -10 | 8.9 |

Brain regions were identified on the Harvard-Oxford Cortical and Subcortical Structural Atlases using the peak-voxel MNI coordinates. ACC = anterior cingulate cortex; MPG = medial prefrontal gyrus; STG = superior temporal gyrus. * Only the first three largest clusters are shown.

**Table S4.** MNI coordinates for cluster peaks associated with inclusion-bearing Layer 5 neighboring neurons in **Figure 2C.**

| **Brain Regions** | **Cluster size (voxels)** | **MNI peak voxel** | **t-values peak voxel** |
|---|---|---|---|
| | | **X Y Z** | |
| **Gray matter atrophy associated with inclusion-bearing Layer 5 neighboring neurons** | | | |
| Right Insula | 1459 | 45 10 2 | 6.2 |
| Right IFG | | 39 28 -4 | 6.0 |
| Right IFG | | 50 18 4 | 5.8 |
| Left Thalamus | 1380 | -2 -4 0 | 6.8 |
| Left Thalamus | | -2 -10 -8 | 6.2 |
| Left Thalamus | | 0 -8 -8 | 6.1 |

Brain regions were identified on the Harvard-Oxford Cortical and Subcortical Structural Atlases using the peak-voxel MNI coordinates. IFG = inferior frontal gyrus; OFC = orbitofrontal cortex

**Table S5.** MNI coordinates for cluster peaks in **Supplementary Figures S2**.

| **Brain Regions** | **Cluster size (voxels)** | **MNI peak voxel** | **t-values peak voxel** |
|---|---|---|---|
| | | **X Y Z** | |
| **White matter atrophy associated with inclusion-bearing VENs and fork cells** | | | |
| Right Forceps Minor | 627 | 16 34 2 | 5.2 |

Brain regions were identified on the John-Hopkins-University White-Matter Tractography Atlas using the peak-voxel MNI coordinates.

**Table S6.** MNI coordinates for cluster peaks in **Figures 3B.**

| Brain Regions | Cluster size (voxels) | MNI peak voxel X Y Z | t-values peak voxel |
|---|---|---|---|
| **Gray matter atrophy associated with inclusion-bearing VENs and fork cells, *C9orf72* non-expansion carriers*** | | | |
| Right Frontal Pole | 10257 | 20 63 -8 | 27.2 |
| Left ACC | | -6 42 8 | 25.5 |
| Right Frontal Pole | | 16 70 -3 | 20.0 |
| Right Insula | 2484 | 52 10 -6 | 17.1 |
| Right PCG | | 60 0 21 | 14.0 |
| Right STG | | 54 2 -2 | 13.6 |
| Right Thalamus | 2164 | 2 -18 14 | 5.2 |
| Right Thalamus | | 3 2 0 | 4.6 |
| Right Thalamus | | 4 3 -9 | 4.2 |

Brain regions were identified on the Harvard-Oxford Cortical and Subcortical Structural Atlases using the peak-voxel MNI coordinates. ACC = anterior cingulate cortex; IFG = inferior frontal gyrus; PCG = precentral gyrus; STG = superior temporal gyrus. * Only the first three largest clusters are shown.

**Table S7.** MNI coordinates for cluster peaks in **Figures 3C**.

| Brain Regions | Cluster size (voxels) | MNI peak voxel X Y Z | t-values peak voxel |
|---|---|---|---|
| **Gray matter atrophy associated with VENs and fork cells TDP-43 pathobiology composite score** | | | |
| Left Thalamus | 1147 | -4 -8 -8 | 8.5 |
| Right Thalamus | | 3 12 -9 | 7.4 |
| Right Thalamus | | 4 -8 -9 | 5.5 |
| Right SFG | 799 | 22 45 36 | 6.5 |
| Right Frontal Pole | | 10 62 14 | 6.5 |
| Right Frontal Pole | | 15 58 22 | 6.3 |
| Left SPG | 623 | -21 -70 50 | 6.3 |
| Left ANG | | -48 -48 48 | 5.6 |
| Left ANG | | 54 2 -2 | 5.1 |

Brain regions were identified on the Harvard-Oxford Cortical and Subcortical Structural Atlases using the peak-voxel MNI coordinates. ACC = anterior cingulate cortex; ANG = angular gyrus; SFG = superior frontal gyrus; SFG = superior parietal gyrus.

### Supplementary Experimental Procedures

#### Specimen and tissue processing

Details regarding specimen and tissue processing can be found in previous work (Nana *et al.*, 2018). Here we provide a summary of the methods used to derive the quantitative neuropathological data re-used in the present study.

#### Autopsy and disease staging

Consent for brain donation was obtained from all subjects or their surrogates in accordance with the Declaration of Helsinki. Postmortem human brain tissue was obtained from the UCSF Neurodegenerative Disease Brain. At autopsy, brains/cerebral hemispheres were either immersion fixed whole and fixed in 10% buffered formalin, or the cerebrum was cut fresh into 1 cm thick coronal slices and fixed for 48-72 hours in 10% buffered formalin. Neuropathological diagnoses were made using previously described histological and immunohistochemical methods (Tartaglia *et al.*, 2010; Kim *et al.*, 2012) following consensus diagnostic criteria (McKeith *et al.*, 2005, 2017; MacKenzie *et al.*, 2010; Mackenzie *et al.*, 2011; Montine *et al.*, 2012). Anatomical disease stage was assessed using an FTD rating scale (Broe *et al.*, 2003); all staging was performed by a single blinded investigator (W.W.S).

#### Specimens and tissue processing

Blocks of the FI were dissected from ~1 cm thick formalin-fixed coronal slabs and stored until processed for sectioning and immunohistochemistry. Blocks were removed from storage in formalin or PBS with 0.02% sodium azide (PBS-Az) and cryoprotected in graded sucrose solutions (10%, 20% and 30% sucrose in PBS-Az then sectioned at 50 μm thickness on a freezing microtome). Every 12th section was Nissl-stained with cresyl violet (FD Neurotech) for neuron type counts and to determine the anatomical boundaries of the right FI, the cortical region of interest.

#### Immunohistochemistry

Three sections were selected from each block for immunohistochemical staining. All sections were coded for rater blinding prior to staining. Free-floating sections were washed (6 × 10 min) in 0.01 M phosphate buffered saline and underwent antigen retrieval in 10 mM citrate buffer pH 6.0 for 2 hours in an 80° C water bath. Endogenous peroxidase activity was quenched with 3% hydrogen peroxide in PBS-Az for 30 min. Sections were then washed (3 × 10 min) in PBS and non-specific staining was blocked by incubating in 5% nonfat milk powder (Carnation) in PBS + 0.25% triton X-100 (PBT) for 1 h. Sections were incubated with an antibody against TDP-43 (Rabbit polyclonal, 1:5000, Proteintech, RRID: AB_615042) on a shaker overnight at room temperature, to determine TDP-43 immunostaining. Following incubation with primary antibody, sections were washed 6 × 10 min in PBS, incubated for 1 h in biotinylated secondary antibody (goat anti-mouse IgG, 1:500 in PBT, Sigma), washed in PBS, and incubated for 1 h in ABC (Vectastain ABC elite kit, Vector Laboratories, Burlingame, CA), washed, then exposed to 0.05% 3,3-diaminobenzidine tetrahydrochloride (Sigma) and 0.01% H2O2 to produce a brown reaction product. The sections were washed, mounted on plus slides, dehydrated through a graded ethanol series, cleared in xylene, and coverslipped with Permaslip. TDP-43 stained sections were counterstained with hematoxylin (Fisher).

**Definition of right FI, image acquisition, and neuron type quantification**

Layer 5 of the right FI was defined as previously described (Kim *et al.*, 2012; Nana *et al.*, 2018). The extent of VENs and fork cells contained in the right FI was determined in cresyl-violet stained slides. Nissl-stained sections from patients using a 60 × objective was used to assess VENs, fork cells, and neighboring neuron loss in right FI Layer 5. Three sections from each subject were assessed while mounted on a motorized stage, using the optical fractionator counting method (StereoInvestigator software, MBF Neuroscience) to obtain apparent local density estimates for each neuronal population. The proportion of each cell type with TDP-43 inclusions or nuclear TDP-43 depletion among total Layer 5 VENs, fork cells, and neighboring neurons was assessed using unbiased cell counting methods. For each section, Layer 5 of the right FI was traced in StereoInvestigator and exported to digital microscope images (Zeiss; 63 × oil, 1.4 NA objective; z-stack interval: 0.4 µm). The numbers of VENs, fork cells, and neighboring neurons in Layer 5 with nuclear TDP-43, TDP-43 depletion and TDP-43 inclusions were then counted using the optical fractionator probe in StereoInvestigator software to obtain the percentage of inclusion-bearing neurons (Nana *et al.*, 2018).

**Inter-rater and intra-rater comparisons**

To obtain fraction rates of inclusion bearing VENs and fork cells, and neighboring neurons, three blinded raters (E.J.K, L.L. and Y.P) performed counting of Nissl-stained sections (Kim *et al.*, 2012; Nana *et al.*, 2018). Intra-rater reliability was assessed independently by each rater every 12 sections. A total of six sections were recounted by each Nissl rater for intra-class correlation analysis. Intra-rater correlation coefficients across the three raters averaged 0.990 (95% CI 0.923-0.999) for VENs, 0.995 (95% CI 0.959-0.999) for fork cells, and 0.987 (95% CI 0.901-0.999) for neighboring neurons.

**Supplementary References**

Broe M, Hodges JR, Schofield E, Shepherd CE, Kril JJ, Halliday GM. Staging disease severity in pathologically confirmed cases of frontotemporal dementia. Neurology 2003; 60: 1005–1011.

Kim EJ, Sidhu M, Gaus SE, Huang EJ, Hof PR, Miller BL, et al. Selective frontoinsular von economo neuron and fork cell loss in early behavioral variant frontotemporal dementia. Cereb Cortex 2012; 22: 251–259.

Mackenzie IRA, Neumann M, Baborie A, Sampathu DM, Du Plessis D, Jaros E, et al. A harmonized classification system for FTLD-TDP pathology. Acta Neuropathol 2011; 122: 111–113.

MacKenzie IRA, Neumann M, Bigio EH, Cairns NJ, Alafuzoff I, Kril J, et al. Nomenclature and nosology for neuropathologic subtypes of frontotemporal lobar degeneration: An update. Acta Neuropathol 2010; 119: 1–4.

McKeith IG, Boeve BF, Dickson DW, Halliday G, Taylor J-P, Weintraub D, et al. Diagnosis and management of dementia with Lewy bodies. Neurology 2017; 89: 88–100.

McKeith IG, Dickson DW, Lowe J, Emre M, O'Brien JT, Feldman H, et al. Diagnosis and management of dementia with Lewy bodies: Third report of the DLB consortium. Neurology 2005; 65: 1863–1872.

Montine TJ, Phelps CH, Beach TG, Bigio EH, Cairns NJ, Dickson DW, et al. National Institute on Aging-Alzheimer's Association guidelines for the neuropathologic assessment of Alzheimer's disease: a practical approach. Acta Neuropathol 2012; 123: 1–11.

Nana AL, Sidhu M, Gaus SE, Hwang J-HL, Li L, Park Y, et al. Neurons selectively targeted in frontotemporal dementia reveal early stage TDP-43 pathobiology. Acta Neuropathol 2018; 137: 27–46.

Tartaglia MC, Sidhu M, Laluz V, Racine C, Rabinovici GD, Creighton K, et al. Sporadic corticobasal syndrome due to FTLD-TDP. Acta Neuropathol 2010; 119: 365–74.
